# A Zika Virus-Like Particle Vaccine Mitigates Early Pregnancy Loss In Rhesus Macaques

**DOI:** 10.1101/2025.10.16.682761

**Authors:** Hannah K. Jaeger, Jessica L. Smith, Caralyn S Labriola, Lydia J. Pung, Olivia L. Hagen, Michael Denton, Rahul J. D’Mello, Christopher J. Parkins, Whitney C. Weber, Samuel Medica, Craig N. Kreklywich, Victor R. DeFilippis, Stephen Bondoc, Kathleen Busman-Sahay, Jenna N. Castro, Gavin Zilverberg, Riely White, Margaret Terry, Aaron M Barber-Axthelm, Michael K. Axthelm, Jeremy Smedley, Matthew T. Aliota, Andrea M. Weiler, Thomas C. Friedrich, Jacob Estes, Terry K. Morgan, Jamie O. Lo, Victoria H.J. Roberts, Daniel N. Streblow, Alec J. Hirsch

**Affiliations:** The Vaccine & Gene Therapy Institute, Oregon Health and Science University, Beaverton, Oregon, USA; Division of Pathobiology & Immunology, Oregon National Primate Research Center, Oregon Health & Science University, Beaverton, Oregon, USA; Division of Reproductive & Developmental Sciences, Oregon National Primate Research Center, Oregon Health & Science University, Beaverton, Oregon, USA; Department of Veterinary and Biomedical Sciences, University of Minnesota, St. Paul, Minnesota, USA; Department of Pathobiological Sciences, School of Veterinary Medicine and Wisconsin National Primate Research Center, University of Wisconsin-Madison, Madison, Wisconsin, USA; Department of Obstetrics & Gynecology, Oregon Health & Science University, Portland, Oregon, USA; Department of Pathology, Oregon Health & Science University, Portland, Oregon, USA

## Abstract

Zika virus (ZIKV) is an arthropod-borne *Orthoflavivirus* that caused a major outbreak in the Americas in 2015-16. In Brazil, up to 46% of ZIKV positive pregnancies resulted in congenital Zika syndrome (CZS). CZS is characterized by a wide range of neurologic birth defects and miscarriage in up to 7.6% of affected pregnancies. With no current licensed ZIKV vaccines, we sought to evaluate a Zika virus-like particle (VLP) vaccine candidate in a rhesus macaque (RM) pregnancy model. VLPs were produced in mammalian cells expressing the pre-membrane-envelope region of the Asian-lineage ZIKV strain PRVABC59, which belongs to the Asian ZIKV lineage that is associated with outbreaks of congenital disease. To evaluate vaccine protection against adverse pregnancy complications, two cohorts of female RM were vaccinated with ZIKV-VLP with adjuvant Alhydrogel (alum) or adjuvant alone prior to mating. At gestational day (GD) 30 (early first trimester), pregnant animals were challenged with ZIKV-DAK 41524, an African-lineage strain shown to induce 1st-trimester fetal demise in 78% (n=11/14 animals) of RM, making it an ideal and stringent model for evaluating ZIKV vaccines. Within the vaccinated cohort, 2 of 3 animals reached the study endpoint of GD 90 with no observed adverse pregnancy outcomes. The third animal experienced pregnancy loss at GD 49 (18 d post infection), although no infectious virus was detected in placental or fetal tissues. In the unvaccinated cohort, two animals had severe adverse events. One animal experienced preterm labor, and another developed early-onset hydrops fetalis with widespread ZIKV-RNA detected via RNAscope and extensive placental damage. These results confirm a significant risk for early pregnancy loss in RM infected with ZIKV-DAK 41524. This model can be further used to understand the complexities of placental immunological features underlying stillbirth and miscarriage following infection. Our findings indicate that this ZIKV-VLP vaccine candidate protected pregnant macaques against fetal demise associated with highly pathogenic ZIKV challenge.

**Author Summary:** Zika virus (ZIKV) infection during pregnancy is associated with pregnancy loss, severe birth defects, including microcephaly and developmental delays, and other subtle neurologic changes. Although vaccine development efforts have been ongoing since the 2015/2016 ZIKV outbreak, no approved vaccine is currently available. Many existing studies have tested vaccines in animal models using strains such as ZIKV-PR that cause mild or moderate pregnancy complications at similar rates to human cases. Using challenge strains that only cause mild, or moderate pregnancy complications makes it difficult to rigorously evaluate vaccine efficacy. In this study, we tested a virus-like particle (VLP) vaccine, a safe and effective method for use during pregnancy as it is replication-deficient and only contains viral structural antigens. We found that the VLP vaccine, when paired with an adjuvant (alum), induced strong antibody responses in mice and controlled viral dissemination following challenge in nonpregnant macaques. To evaluate the protective efficacy of the ZIKV-VLP vaccine against in utero infection and disease, we used a stringent model of ZIKV infection during early pregnancy in rhesus macaques that is associated with high rates of fetal demise. In pregnant macaques, the vaccine reduced maternal viremia, limited viral spread to fetal and placental tissues, and conferred protection against virus-mediated fetal demise and placental damage. This is the first study to evaluate a VLP-based vaccine in a consistent pregnancy loss model of ZIKV infection. These findings support the continued development of VLP vaccines as a safe and effective strategy for protecting pregnant individuals and their developing fetuses from ZIKV.

## 1. Introduction

Zika virus (ZIKV), a mosquito-borne orthoflavivirus, remains a critical global public health concern due to its ability to cause birth defects and adverse pregnancy outcomes, including miscarriage, intrauterine growth restriction (IUGR), and stillbirth (1–10). ZIKV was first isolated in Uganda in 1947 and has since diverged into two distinct lineages: African and Asian (11–13). The Asian lineage circulated in mainland Southeast Asia before being introduced into the Pacific region, which eventually led to an explosive outbreak in South America in 2015 (14). Brazil had the most cases during this outbreak, with up to 46% of ZIKV+ pregnancies developing adverse outcomes, including a miscarriage rate of 7.6% (1). Since the outbreak, several fetal neurological defects have been associated with ZIKV infection during pregnancy, ranging from subtle to severe, most notably microcephaly. Collectively these defects are known as Congenital Zika Virus Syndrome (CZS) (2, 9, 15, 16).

To better understand the mechanisms of ZIKV-induced birth defects and evaluate potential countermeasures, several studies have utilized nonhuman primate (NHP) models—particularly rhesus and cynomolgus macaques—as a translational platform. NHP pregnancy models offer many advantages, including the similarity of the placenta structure, mode of placentation, gestational time, fetal brain development – especially prefrontal cortex expansion, and immune responses relative to humans (17, 18). Importantly, vertical transmission, placental pathology, and fetal neurologic outcomes observed in macaques also closely mirror those seen in humans (19). Most NHP studies to date have utilized Asian-lineage ZIKV strains, including isolates from Puerto Rico, Brazil, French Polynesia, and Cambodia. First-trimester infection produces the most severe outcomes, with a meta-analysis reporting a pregnancy loss rate of ∼26% in macaques infected with ZIKV strain PRVABC59/2015 (ZIKV-PR) at gestational day 30 (20). These models have been instrumental in characterizing pathogenesis and host immune responses but result in only moderate rates of fetal demise.

Recent efforts utilizing the African-lineage ZIKV strain ZIKV/*Aedes africanus*/SEN/DAK-AR 41524/1984 (ZIKV-DAK) have demonstrated consistent severe adverse fetal outcomes in NHPs, which is desirable for testing countermeasures, enabling appropriately powered experiments with smaller group sizes. In NHPs, infection with ZIKV-DAK at GD30 results in rapid and severe fetal demise in up to 78% of pregnancies (21, 22) and was observed as early as 6 days post-infection (dpi) in fetal placenta tissues and membranes (23). As such, it serves as a rigorous, high-penetrance model for evaluating the protective efficacy of ZIKV vaccine candidates in the context of early pregnancy infections.

Despite the need for interventions that protect against adverse outcomes, the ZIKV vaccine pipeline has faced significant obstacles. Several vaccines have been developed for ZIKV infection with a select few making it to clinical trials (24). As of October 2025, 20 phase 1 clinical trials have been completed using vaccine platforms such as live attenuated, inactivated whole virion, mRNA, DNA, virus-vectored, and recombinant peptides (24–31). Three candidates advanced to phase 2 clinical trials but were withdrawn (Takeda TAK-426, NCT05469802), or discontinued due to a lack of funding (Moderna mRNA-1893, NCT04917861) (32), and one was completed early without assessment of efficacy assessment (National Institute of Allergy and Infectious Diseases VRC5283, NCT03110770). Since the peak of the 2015–2016 outbreak, ZIKV transmission has decreased, making it more challenging for clinical trials to evaluate vaccine efficacy, even in areas where ZIKV may be endemic; this is a significant barrier to advancing vaccine candidates (33).

A notable advancement in vaccine testing is the development of the ZIKV human challenge model (34). However, in general, these models and clinical trials have excluded pregnant individuals due to ethical and safety concerns. Moreover, many platforms, including replicating viral vectors and live-attenuated viruses, pose safety risks during pregnancy due to the potential for viral replication, making them unsuitable for use in vulnerable populations (35–38). Virus-like particle (VLP)-based vaccines are a promising platform for the development of a ZIKV vaccine due to improved safety profiles for use during pregnancy. Expression of the pre-membrane/envelope (prME) region of the ZIKV genome results in the self-assembly and secretion of VLPs that mimic the original structure of the virus. Since these particles do not contain viral genomic material or replication machinery, they are nonreplicating and safer than attenuated vaccines, which pose potential health risks to vulnerable populations such as neonates, pregnant women, elderly individuals, as well as those living with compromised immune systems. VLPs have been shown to induce effective and durable humoral responses in both mosquito-borne and tick-borne flaviviruses when adjuvanted to enhance immunogenicity (39–41).

In this study, we evaluated the safety, immunogenicity, and protective capacity of a ZIKV-VLP vaccine adjuvanted with alum in both mice and rhesus macaques. In nonpregnant animals, we characterized antibody responses, neutralization capacity, and tissue dissemination following viral challenge. We found similar immune responses between males and females, with the highest immunogenicity seen in the cohort receiving the adjuvanted VLP vaccine. To assess efficacy in pregnancy, we utilized the ZIKV-DAK strain early-pregnancy loss model by challenging rhesus macaques at 30 days gestation (GD30) and monitoring outcomes over 60 days. Comprehensive assessments included longitudinal viremia and immune profiling through routine blood draws, alongside serial non-invasive ultrasound (US) evaluations to track fetal growth, uteroplacental blood flow, and amniotic fluid index. At necropsy, fetal, placental, and maternal tissues were analyzed for viral burden, histopathology, and infectious virus recovery. This study represents the first application of a VLP-based vaccine in a stringent, high-penetrance NHP model of ZIKV-induced pregnancy loss. Our results provide novel insight into the ability of VLP vaccination to reduce maternal viremia, prevent vertical transmission, and mitigate adverse fetal outcomes, which offers a valuable step forward in the development of safe, effective ZIKV vaccines for use in pregnancy.

## 2. Results

### 2.1 ZIKV-VLP adjuvanted with alum elicit robust immune responses in mice and rhesus macaques

Zika virus-like particles (VLP) were collected from culture supernatants of HEK293 or Vero cells expressing the ZIKV prM-E coding region, followed by purification and concentration over a 20% sorbitol cushion. VLP were analyzed by SDS-PAGE and western blot and quantified relative to a titered ZIKV-PR control **(S Figure 1)**, yielding an estimated concentration of 5×10⁶ focus forming equivalents (FFE)/μL.

**Figure 1.**
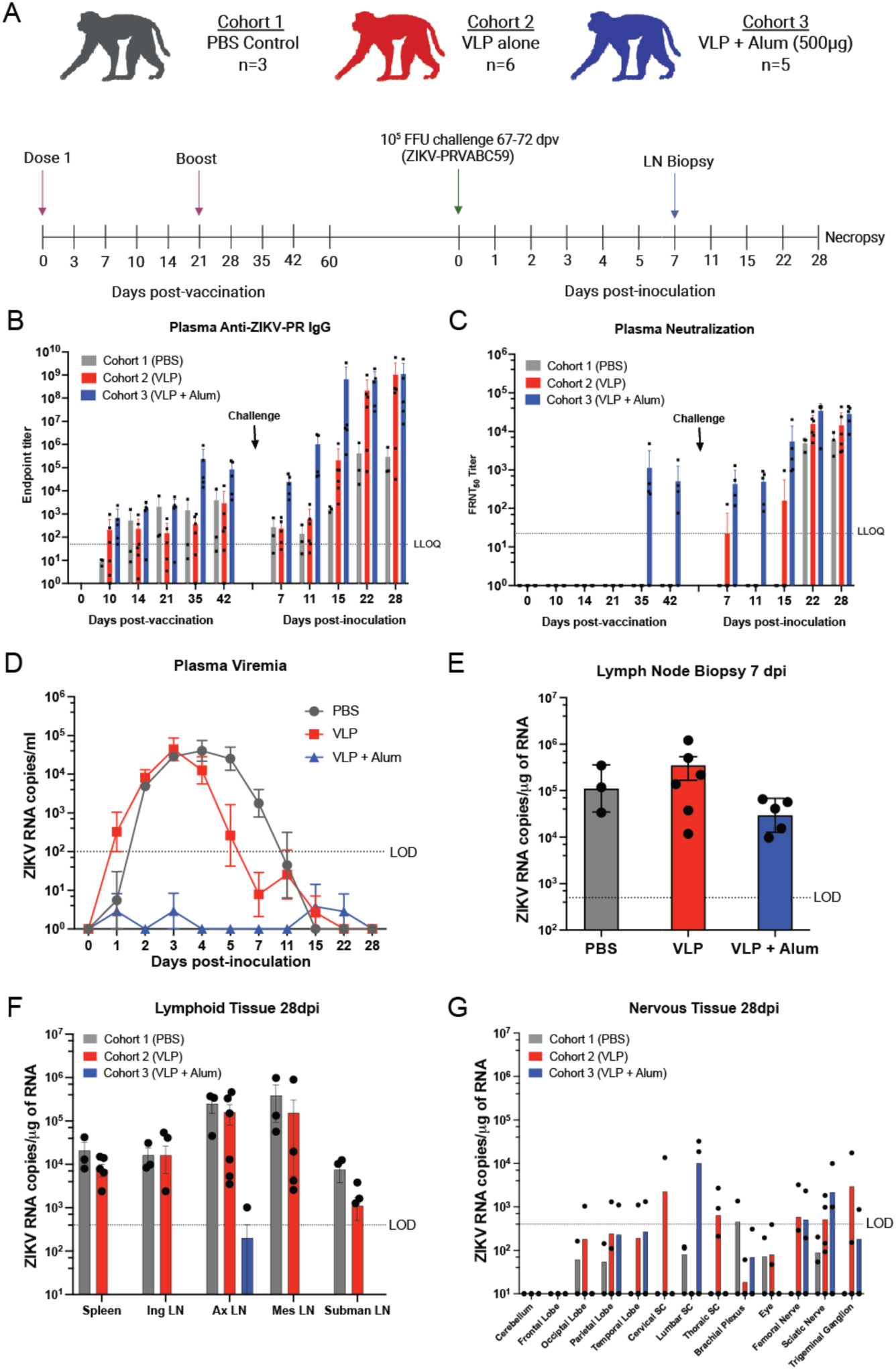
**ZIK-VLP vaccine adjuvanted with alum induces ZIKV- neutralizing antibodies and limits ZIKV replication and spread in nonpregnant macaques. *A***, Study schematic for cohorts 1–3. Non- pregnant rhesus macaques (average ages: 9.6, 10.9, and 11.4 years for cohorts 1–3, respectively) were immunized subcutaneously (s.q.) in the right arm with either PBS (control), ZIKV-VLP alone (10^5^ FEE), or 10^5^ FEE ZIKV-VLP adjuvanted with 500 μg alhydrogel (alum). Animals received a prime and boost 21 days apart, followed by challenge with ZIKV strain PRVABC59 (ZIKV-PR). Challenge was performed with a total of 10⁵ FFU administered via five s.q. injections per arm (10⁴ FFU per site) to mimic mosquito bites. Blood was collected at designated timepoints to assess plasma viremia and immune responses. Schematic was created in BioRender. Jaeger, H. (2025) https://BioRender.com/wnxj8so. ***B***, ZIKV- binding IgG titers were measured in plasma via ELISA. The lower limit of quantification (LLOQ) was 1:50 (dashed line). ***C***, ZIKV-neutralizing antibody responses were assessed using focus reduction neutralization tests (FRNT₅₀) with heat-inactivated plasma. Titers were calculated using nonlinear regression. The LLOQ was a 1:50 plasma dilution. ***D***, Plasma viremia was quantified via RT- qPCR. Data represent the mean of three technical replicates. The limit of detection (LOD) was 100 ZIKV RNA copies/mL, as determined by standard curve. ***E***, Axillary lymph node (LN) biopsies were collected under ultrasound guidance at day 7 post- infection and viral RNA was quantified per 1 μg of tissue. ***F, G*** Viral loads at necropsy (28 dpi) were detected via one-step RT- qPCR in triplicate. The LOD was 100 copies of vRNA/µg.

The immunogenicity of ZIKV-VLPs was first assessed in a mouse model. C57BL/6 mice were primed by intramuscular (i.m.) injection with 1×10^5^ FFE alone (n=4) or adjuvanted with 150 μg alhydrogel (alum) (n=4). At 21 days post-prime, mice were boosted by i.m. injection with the same vaccine formulations and dosage. Serum was collected at 6 weeks post-prime for analysis by ELISA and neutralization titration assays **(S Figure 2A)**. While mice receiving the unadjuvanted vaccine did not demonstrate detectable antibody responses, the adjuvanted VLP vaccine elicited detectable total IgG responses against whole viral antigens with a dilution titer of approximately 3×10^4^. Similar IgG1 and slightly lower IgG2c titers (4×10^3^) were detected in serum from this group **(S Figure 2B-D).** Additionally, the Alum adjuvanted VLP vaccine elicited neutralizing antibody titers of 1×10^3^, detected by 50% focus reduction neutralization tests (FRNT_50_) against ZIVK-PR **(S Figure 2E).** These results demonstrate that vaccination with an adjuvanted formulation of ZIKV-VLPs induces a potent neutralizing immune response in mice, providing a rationale to advance into the rhesus macaque model.

**Figure 2.**
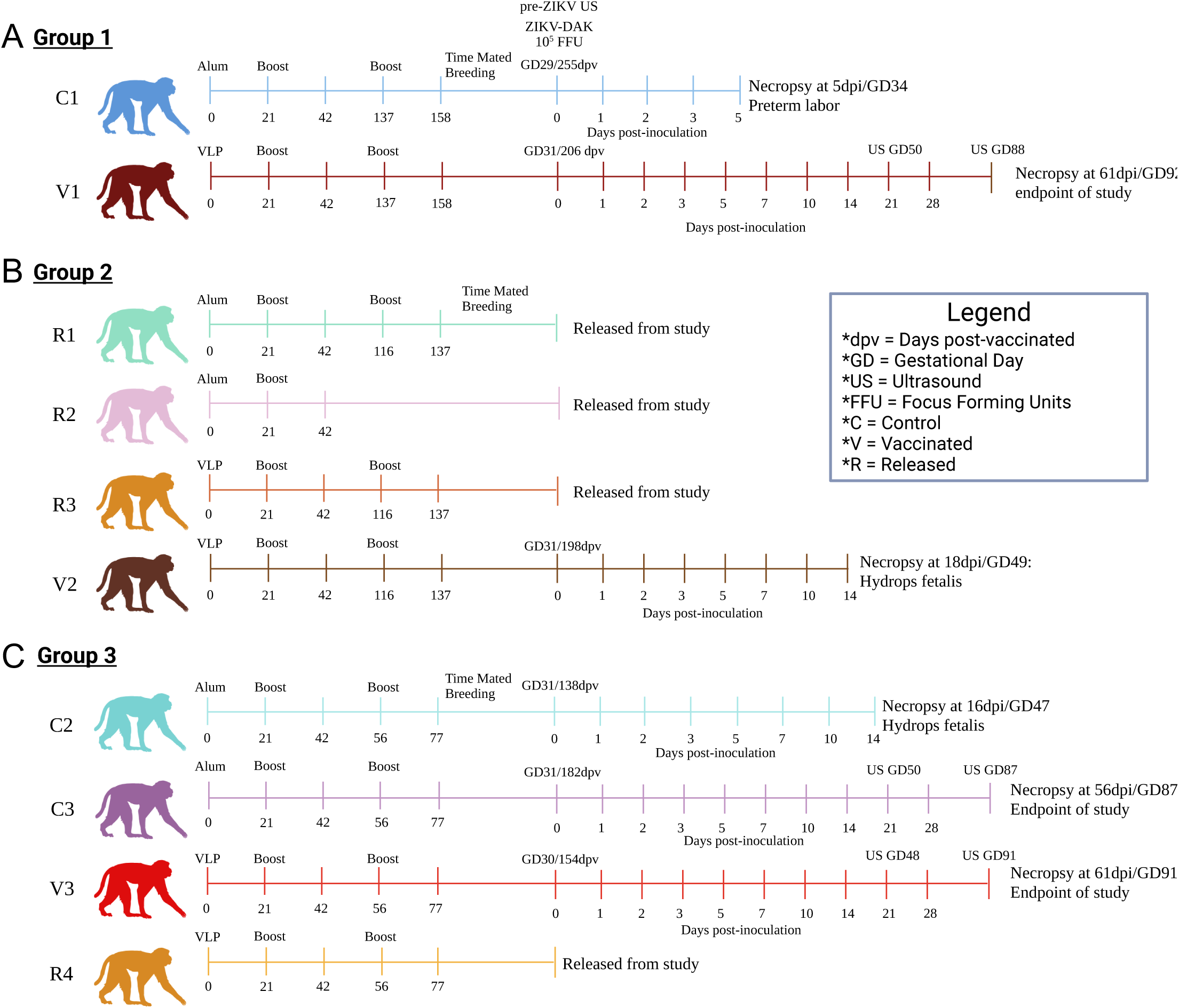
ZIKV-VLP vaccination in pregnant macaques: study design and fetal outcomes. *A-C*, Female rhesus macaques (average age: 15 years) were immunized s.q. in the right arm with either 500 μg alum alone (n=5) or 10⁵ FFE of ZIKV-VLP adjuvanted with 500 μg alum (n=5) for a total of three doses. Animals then were placed into time-mated breeding (TMB) and were subsequently challenged with ZIKV strain Dakar 41524 (ZIKV-DAK). Animals were challenged with 10⁵ FFU, administered across ten total s.q. sites with 5 sites per arm (10⁴ FFU per site) to mimic multiple mosquito bites. Blood samples were collected at designated timepoints to assess serum viremia and immune cell activation. Animals were also monitored daily for signs of viremia and weight loss and underwent non-sedated ultrasound assessments every other day to monitor fetal viability. Females were divided into three groups, resulting in different average days post-vaccinated to challenge: ***A,*** Group 1 (230 dpv) ***B***, Group 2 (198 dpv) and ***C,*** Group 3 (158 dpv). *In alum-only control group (C), one fetus was euthanized following preterm labor with bradycardia (C1), and another experienced fetal demise due to features consistent with hydrops fetalis (14–16 dpi; C2). In the VLP + Alum group (V), one case of hydrops fetalis met humane endpoint criteria at 18 dpi (V2). Animals that did not become pregnant were released from the study and are denoted by R. Image created in BioRender. Jaeger, H. (2025) https://BioRender.com/dqz39uj

We next evaluated ZIKV-VLP immunogenicity in three cohorts of adult rhesus macaques (*Macaca mulatta*, RM) consisting of both males and nonpregnant females that were vaccinated by intramuscular injection (i.m.) **(Figure 1A)**. Cohort 1 received a PBS vehicle control (n=3). Cohorts 2 and 3 received priming doses of 5×10^6^ FFE of ZIKV-VLP (n=6) and 5×10^6^ FFE of ZIKV-VLP adjuvanted with 500 µg of alum (n=5), respectively. All animals received a homologous vaccine boost at 21 days post-prime. Blood samples were collected throughout the study at the indicated times **(Figure 1A)**. Between 67 and 72 days post-prime, all groups were challenged with 10^5^ FFU ZIKV-PR s.q. Lymph node biopsies were collected at day 7, and animals were necropsied at 28 days post-challenge **(Figure 1A)**.

Plasma samples collected pre- and post-vaccination were assessed for ZIKV-reactive IgG by ELISA using plates coated with virus stocks **(Figure 1B).** Anti-ZIKV IgG responses were most robust in the adjuvanted VLP group (Cohort 3) starting at 28-35 days post-vaccination (dpv) with peak dilution titers equivalent to approximately 1×10^5^. By 2 weeks post-ZIKV challenge, occurring at day 81 dpv, all three groups exhibited increased levels of ZIKV-reactive antibodies. Cohort 3 animals mounted the most rapid and potent antibody response **(Figure 1B).** Focus forming reduction neutralization activity for pre-challenge serum (42 days post-vaccination) was only detected for Cohort 3 (FRNT_50_ average value of 1×10^3^) **(Figure 1C)**. However, following challenge, plasma from all animals exhibited neutralization activity that increased with time, with Cohort 3 retaining the highest levels, peaking at dilutions of approximately 2×10^4^ **(Figure 1C)**. These findings demonstrate that adjuvanted ZIKV-VLP vaccination elicits ZIKV-specific antibody responses and neutralizing activity in both mice and rhesus macaques.

### 2.2 ZIKV-VLP reduces plasma viral loads and tissue dissemination in nonpregnant rhesus macaques

To evaluate the protective efficacy of ZIKV-VLP vaccination against viral replication and tissue spread, viral RNA was quantified in plasma at multiple timepoints, from lymph node biopsies at 7 dpi, and tissues collected at necropsy at 28 dpi. Viral RNA was detected in the plasma of Cohort 1 (PBS control, grey) from days 1 to 5 post-challenge, with peak levels observed around day 3 post-inoculation **(Figure 1D)**. Cohort 2 (VLP vaccinated, red) displayed similar viremia kinetics, although the levels of viral RNA were reduced by almost 2 logs at day 5 and 7, which suggests this group experienced a shortened duration of viremia. We further evaluated this by analyzing the area under the curve for each cohort (PBS: 20.71±2.077; VLP: 17.98±3.577; VLP/alum: 0±0) and found the AUC of VLP adjuvanted with alum was significantly reduced when compared to PBS control or VLP alone (p<0.0001). Notably, no viremia above the limit of detection was detected in Cohort 3, demonstrating that vaccination with adjuvanted ZIKV-VLP is sufficient to control systemic spread of infectious virus **(Figure 1D)**. Viral RNA was detected in draining lymph nodes (axillary lymph nodes) on day 7 post-challenge by RT-qPCR in all groups **(Figure 1E)**, additionally the rise in neutralizing antibodies **(Figure 1C)** confirms that the protection observed in Cohort 3 was not due to failed inoculation but indeed a reflection of the protective effects of this vaccination strategy. Prior studies from our group and others demonstrated that ZIKV RNA can persist in lymphoid and nervous system tissues in rhesus macaques for up to 40 days post- inoculation (42–44). To assess the impact of vaccination on viral persistence, we performed RT-qPCR on lymphoid and nervous system tissue samples collected during necropsy at day 28 post-challenge **(Figure 1F and G)**. Persistent ZIKV RNA was detected in the lymph node tissues from both Cohorts 1 and 2, but not in the group that was vaccinated with the adjuvanted ZIKV-VLP, except for one animal with a low, but detectable, level of viral RNA in the axillary lymph node **(Figure 1F)**. In addition, a range of nervous tissue samples from all three groups were positive for viral RNA **(Figure 1G)**, with no correlation between viral detection and vaccination status. Overall, these findings indicate that vaccination with adjuvanted ZIKV-VLPs effectively limits viremia and systemic ZIKV replication as well as persistence in lymphoid compartments, supporting the potential of this platform as a protective ZIKV vaccine.

### 2.3 ZIKV-VLP provided partial protection against an early-pregnancy loss in rhesus macaques

To assess the prophylactic efficacy of the ZIKV-VLP vaccine during pregnancy, five nonpregnant female RM were vaccinated with 5×10^6^ FFE VLPs adjuvanted with 500 µg of alum. Five additional nonpregnant time-mated breeding (TMB) animals that received alum alone served as controls. To ensure appropriate age-matched comparisons, the animals were divided into three staggered infection groups **(Figure 2A-C, S Table 1)**. Dams were primed (i.m.) and boosted 21 days after priming vaccination. Blood samples were taken for serologic analyses and tested for IgG levels at 42 days post prime **(Figure 3A)**. Titers were determined to be lower than the nonpregnant cohort **(Figure 1B)**, and the animals were boosted a second time to achieve comparable ELISA titers before enrollment in TMB program. Dams were then enrolled in TMB **(Figure 2A-C),** and pregnancies were confirmed via ultrasound (US). Four dams failed to conceive, and these animals were released from the study (R1- 4). Dams with US-confirmed pregnancies were challenged by subcutaneous (s.q.) injection in the arms with 10^5^ FFU of ZIKV strain Dakar 41524 (ZIKV-DAK) at GD30 +/- 1 day, a timepoint previously shown to yield high fetal loss rates (21). Fetal viability was monitored with frequent, non-sedated US to identify early pregnancy loss, typically observed between 12–20 days post-infection (dpi) in this infection model (21). In our study, three animals (C1, C2, and V2) experienced clinical signs of early fetal compromise or demise in utero within this timeframe, while the remaining three animals (C3, V1, and V3) carried their pregnancies to the study endpoint (GD87-92).

**Figure 3.**
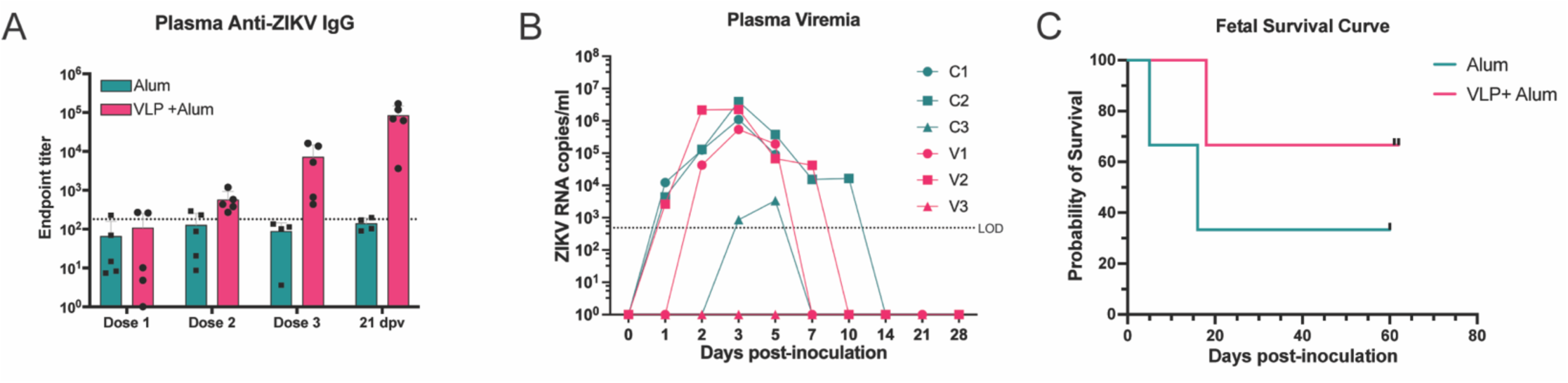
**Dynamics of ZIKV-Specific Antibodies, Maternal Viremia, and Fetal Survival. *A,*** ZIKV-binding IgG antibody titers were quantified in maternal macaque plasma at the indicated timepoints by ELISA. The dashed line represents the average baseline (day 0) titer, determined to reflect background/nonspecific binding. The lower limit of quantification (LLOQ) was a 1:20 dilution. ***B,*** Maternal plasma viremia was measured by RT-qPCR, with each data point representing the average of three technical replicates. The limit of detection (LOD) was 500 copies ZIKV RNA/mL plasma; samples below the LOD are plotted at 0 copies/mL. ***C,*** Kaplan-Meier survival curve of fetal outcomes stratified by treatment group.

**Table 1.**
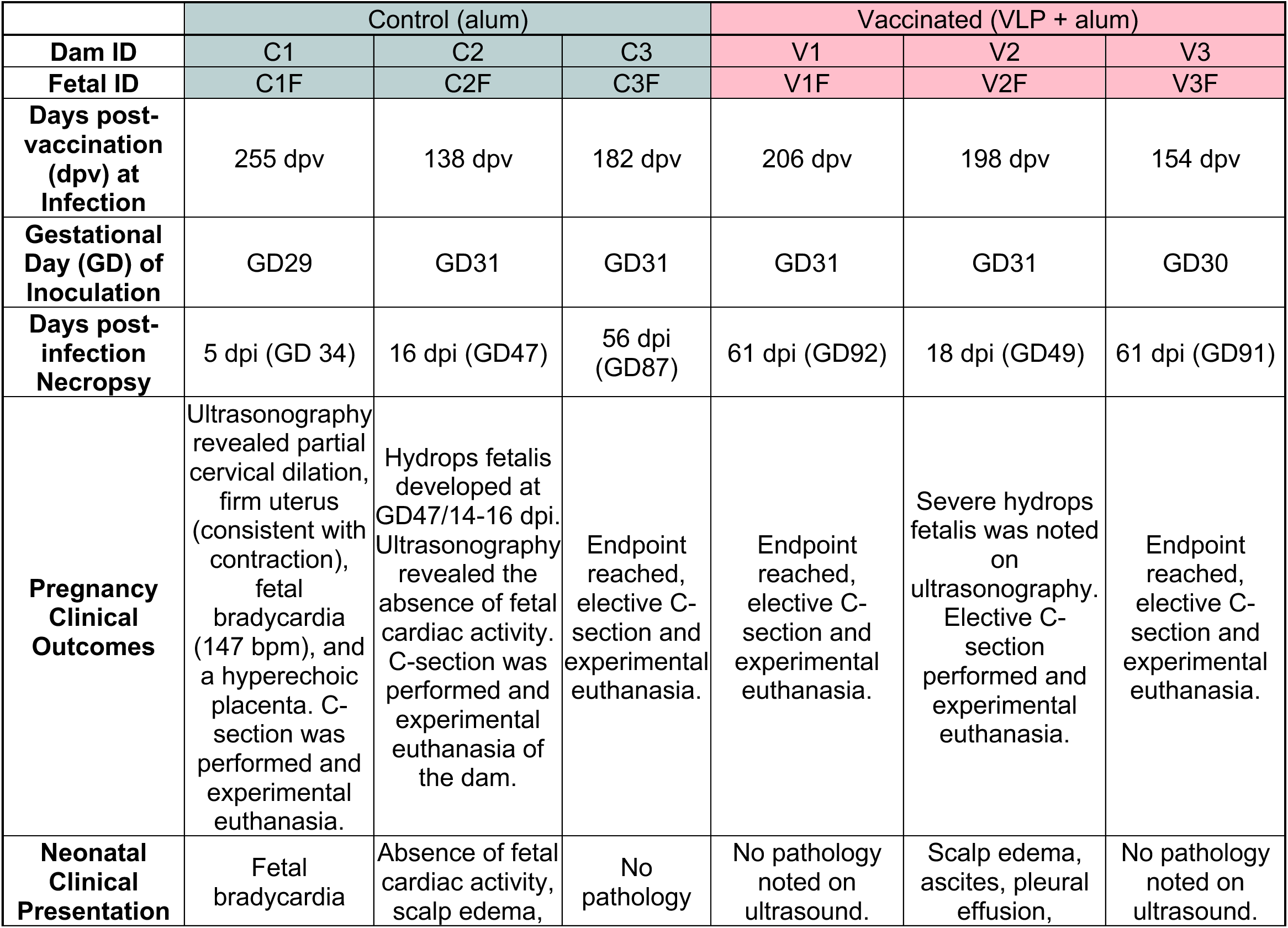
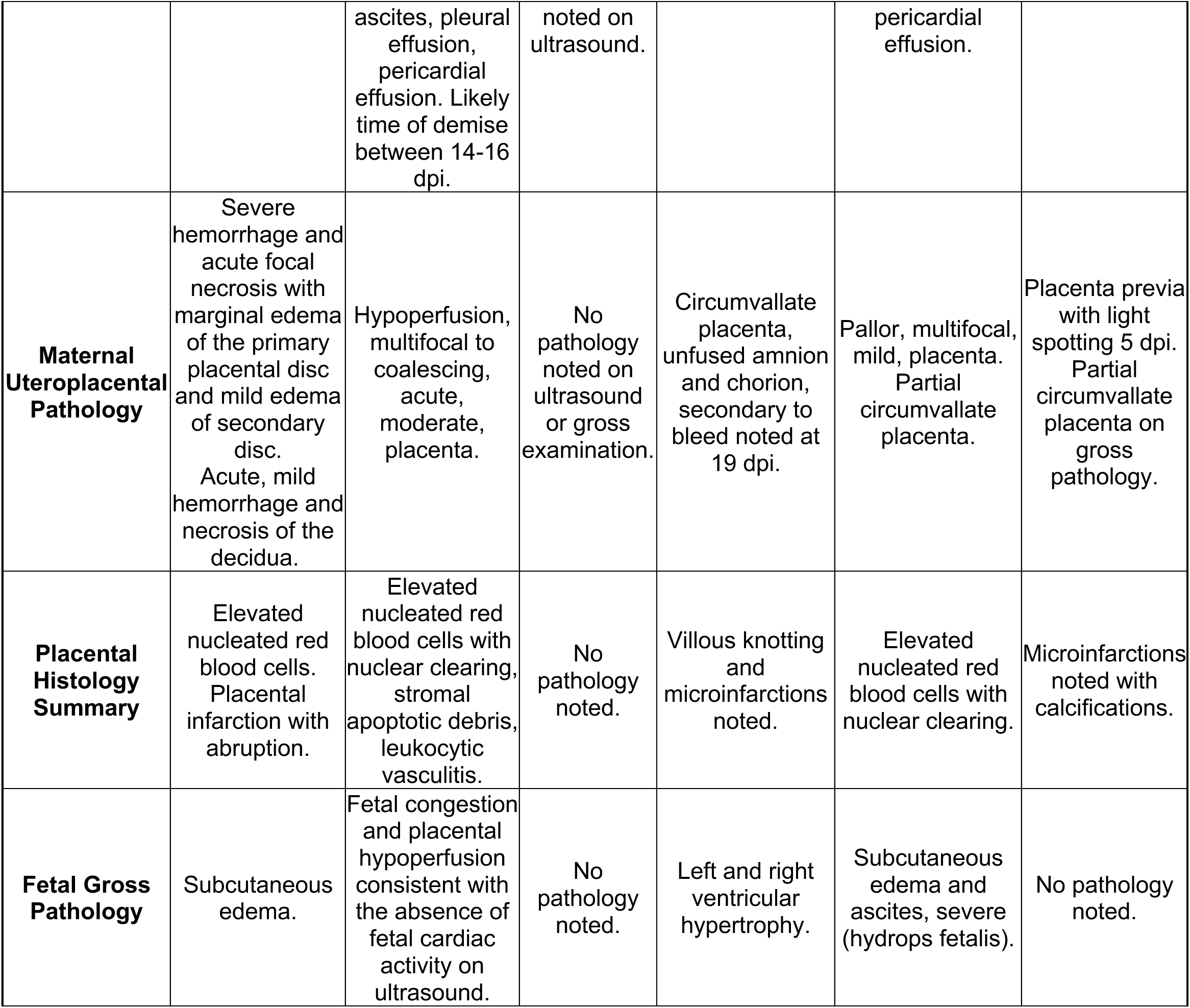
Summary of clinical and necropsy findings. Overview of key clinical observations and necropsy results for each animal included in the study. Clinical findings may include changes in behavior, appetite, or activity, while necropsy findings document gross pathological changes observed at the time of tissue collection. Data are presented for maternal and fetal (F) assessments where available.

To determine if maternal viremia was associated with fetal demise, RNA was extracted from plasma and quantified by ZIKV-specific quantitative PCR. Pregnant animals that experienced fetal demise (C1, C2, V2) showed earlier maternal onset of viremia (1 dpi), with peak levels by 3 dpi and sustained high titers until shortly before fetal loss **(Figure 3B)**. In contrast, animals that maintained pregnancy to endpoint (C3, V1, V3) exhibited delayed or undetectable levels of viral RNA (V3 had no detectable viremia, and C3 had a modest peak at 5 dpi). V1 had detectable viremia; however, the onset was delayed by one day and the duration and amplitude of virus detection were reduced in comparison to the early demise cases. Kaplan-Meier survival curve of fetal outcomes was stratified by treatment group, showing a rate of demise of 66% in control animals and a reduced rate of 33% in vaccinated animals **(Figure 3C)**. Based on these outcomes, animals were stratified into two groups: the early demise cases (C1, C2, and V2) and endpoint cases (C3, V1, and V3).

Activation of blood monocyte/macrophage and dendritic cells (DC) has been a reproducible marker of ZIKV infection and corresponds with viremia (42, 43, 45). PBMCs collected following infection were stained for macrophage/DC subset markers and CD169 **(S Fig 3)**. Activation of macrophage subsets and DCs was observed for all animals with the levels corresponding to the degree of viremia. The control group demonstrated earlier activation and greater frequency of CD169+ classical and intermediate monocytes than the vaccinated cohort **(Figure 4A-D)**. A similar trend was observed in T-cell-stained subsets, with CD8+ T-effector memory cells having the highest granzyme B curves in the early demise cases (C2, V2) **(S Fig 4,5)**. While ZIKV binding and neutralizing antibodies increased over baseline more rapidly, by 14 days post-inoculation and at subsequent timepoints, the levels were generally normalized across both groups **(Figure 5A-B)**. It is important to note that neutralizing antibodies for two of the vaccinated animals (V1, V2) fell below the level of detection prior to challenge, which could be due to the extended time post vaccination in our pregnancy cohorts. Taken together, these results demonstrate that, while adjuvanted ZIKV-VLP vaccination confers partial protection, it does not absolutely prevent early pregnancy loss.

**Figure 4.**
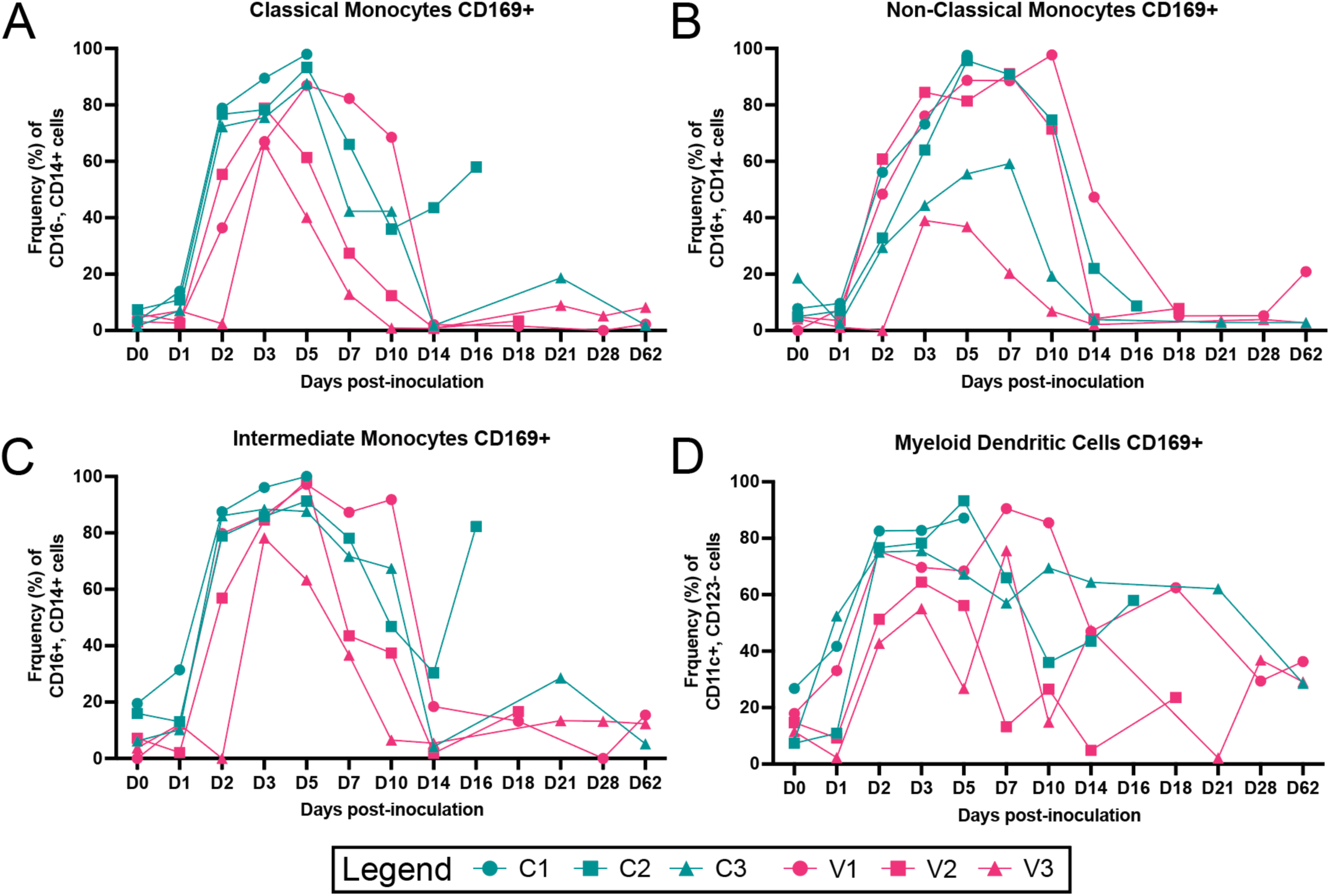
Longitudinal peripheral blood innate immune cell phenotype and activation. Rhesus macaque PBMC isolated at the indicated timepoints were stained with antibodies directed at specific cellular markers and analyzed for cell phenotype using flow cytometry. Changes in the longitudinal frequency of both total and activated (CD169+) classical monocytes ***(A)***, non-classical monocytes ***(B)***, and intermediate monocytes ***(C)*** were quantified. Dendritic cells (CD16-/CD14-) were separated into myeloid dendritic cells ***(D)***, and plasmacytoid dendritic cells (not shown) were quantified as total and activated (CD169+). Lines represent individual animals referenced in the legend with Alum (teal) and VLP-Alum (pink).

**Figure 5.**
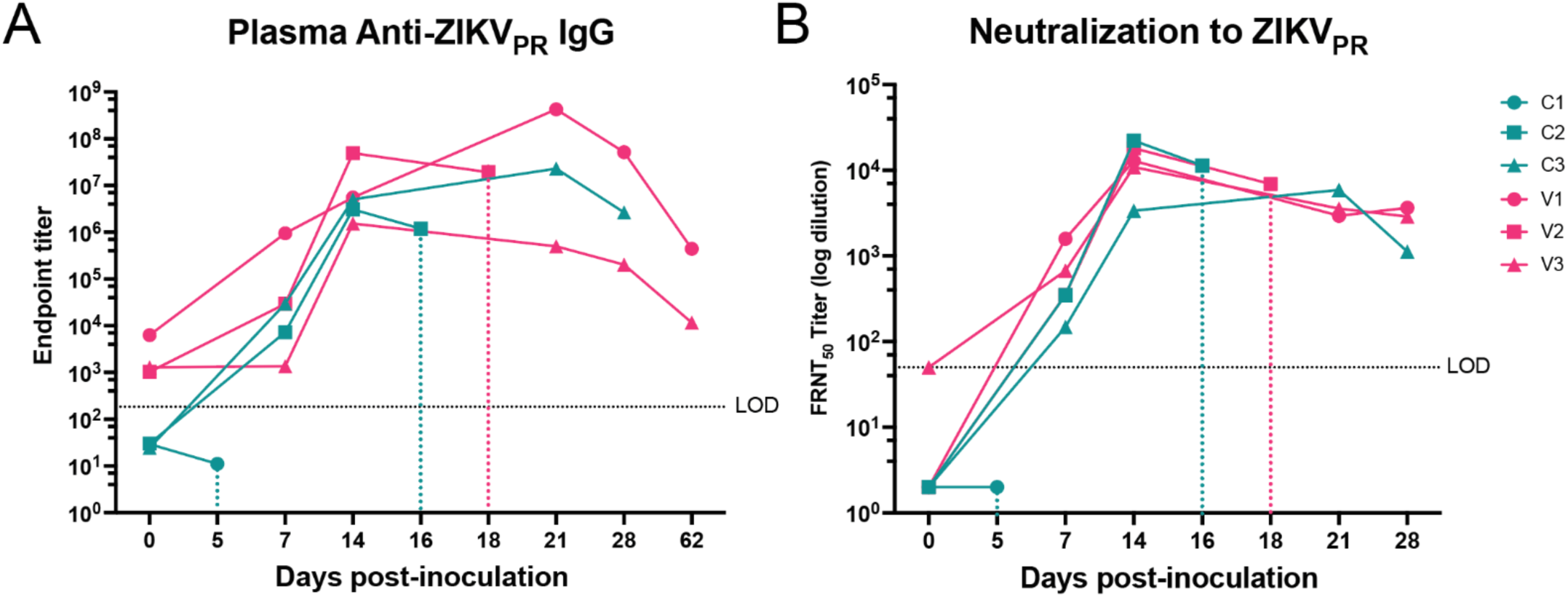
**Adaptive humoral responses show differences in early timepoints post-ZIKV inoculation. *A***, ZIKV- binding IgG antibody titers were quantified in maternal macaque plasma at the indicated timepoints by ELISA. The dashed line represents the average baseline (day 0) titer for all animals (Figure 3A), determined to reflect background/nonspecific binding. The limit of detection (LOD) was a 1:20 dilution. ***B***, the longitudinal development of ZIKV-neutralizing antibodies was quantified in ZIKV neutralization assays using the ZIKV-PR strain as the infectious virus and heat-inactivated macaque plasma at indicated timepoints post-inoculation. The 50% focus reduction neutralization titers (FRNT_50_) were determined using non-linear regression. The LOD was a 1:50 plasma dilution indicated by the dashed lines (black). Vertical dashed lines indicate the timing of demise of each fetus.

### 2.4 Early pregnancy loss clinical features suggest hydrops fetalis

To investigate the potential causes of fetal demise, dams and fetuses were monitored for clinical signs, ultrasound changes, and tissue pathology findings upon necropsy **(Table 1)**. Control 1 fetus (C1F) died on day 5 post-inoculation. The pregnancy was complicated by fetal bradycardia and maternal spotting and uterine contractions concerning for preterm labor. Ultrasonography revealed partial cervical dilation and a firm uterus consistent with contractions and a hyperechoic placenta, which is atypical at this gestational age and is suggestive of abnormal placental pathology. A cesarean delivery was performed, and postnatal examination revealed diffuse subcutaneous edema in the fetus. Control animal C2F also died early at 14-16 dpi. Fetal cardiac activity was normal at 14 dpi but absent 48 hours later, with the presence of fetal pericardial effusion and scalp edema, prompting surgical delivery. Necropsy revealed findings consistent with hydrops fetalis, a condition with fluid accumulation in at least two fetal compartments (i.e., the abdomen, chest, or skin) that is associated with increased mortality, as the cause of fetal demise for C2F. There are few reported cases of ZIKV associated with nonimmune hydrops fetalis (NIHF), suggesting maternal antibodies are not the mechanism for fetal red blood cell destruction (46, 47). NIHF secondary to infection in humans has a time course of about 7-10 days from diagnosis to demise and commonly occurs in 2^nd^ trimester. (48). In our rhesus macaque study, the timing appears to be accelerated, with clinical features detected by US 48 hrs. prior to demise. Animal C3F had no complications throughout pregnancy and was carried to the study endpoint at GD90. These data confirm ZIKV-DAK infection in early pregnancy carries a high risk of fetal demise and is a stringent challenge model for evaluating countermeasures against CZS.

Among vaccinated animals, a similar case of hydrops fetalis was seen at day 16 post-inoculation. The fetus (V2F) developed features consistent with hydrops fetalis, including scalp edema **(Figure 6A)**, ascites **(Figure 6B)** and pleural effusion **(Figure 6C)**, ultimately resulting in end-organ failure. A cesarean delivery was performed, and the fetus and dam were humanely euthanized once the fetus developed signs of severe hydrops at 18 dpi. Diffuse generalized subcutaneous edema was noted on US in both C2F and V2F **(Figure 6D)** and confirmed on necropsy **(Figure 6E)**. Additionally, hepatomegaly was observed at necropsy, indicative of end-organ failure. Vaccinated dams V1 and V3 carried their pregnancies to endpoint at GD90 but had noted changes seen on US. V1 exhibited amniotic and chorionic membrane separation beginning at 19 dpi, concerning for a bleed that continued throughout the pregnancy **(Figure 6F)**. A circumvallate placenta was also noted, which is associated with placental dysfunction and increased risk for pregnancy loss **(Figure 6G)** (49). Dam V3 experienced intermittent spotting throughout early pregnancy, likely secondary to the presence of a placenta previa, which was seen on US before inoculation **(S Figure 6)**.

**Figure 6.**
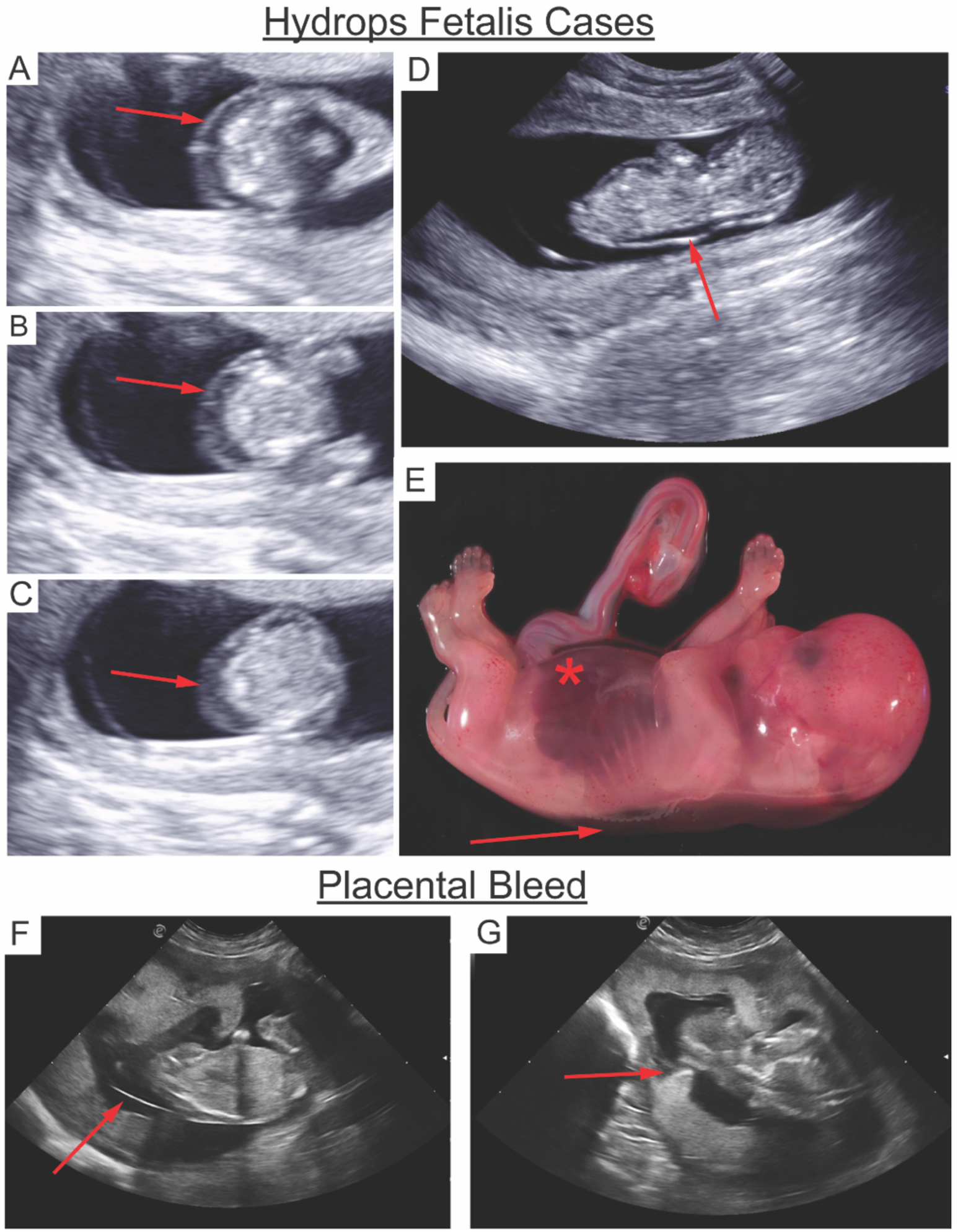
**Representative ultrasound and necropsy findings in ZIKV-infected pregnancies. *A-D,*** Ultrasound images from cases C2F and V2F (gestational days 46–48) show features of hydrops fetalis, including scalp edema **(*A*)**, ascites **(*B*)**, pleural effusion and cardiomegaly ***(C)***, and generalized subcutaneous edema spanning the entire fetus ***(D)***. ***E,*** Necropsy confirmed these findings, with the red asterisk denoting hepatosplenomegaly and the red arrow highlighting subcutaneous edema and spinal cord tenting. ***F,*** A large placental bleed with chorionic and amniotic membrane separation was observed in case V1 at 35 days post-infection. ***G,*** The placental edge folded over consistent with a circumvallate placenta was also noted in this animal. All ultrasound images were obtained by board-certified maternal-fetal medicine (MFM) physician J.O.L.

### 2.5 Fetal growth dynamics in endpoint cases indicate healthy growth dynamics

Maternal weight was monitored throughout infection for each dam and expressed as a percentage of their baseline weight. All three control animals exhibited weight loss during the first 2–3 weeks post-inoculation, with the most pronounced reductions observed in those that experienced early fetal demise **(Figure 7A)**. While overall maternal clinical assessments were unremarkable, this weight loss may represent an early physiologic indicator of adverse pregnancy outcome. To investigate any impacts on fetal growth, US analysis was used to measure fetal growth dynamics, including biparietal diameter, head circumference, abdominal circumference, and femur length. Fetal growth restriction has been observed in animal models and human ZIKV cases, which is known to lead to a reduction in the growth dynamics (1, 7, 8, 50). Notably, in ZIKV cases, fetal growth restriction appears to have femur length sparing, with more evident changes seen in abdominal and head circumference (8). Animal studies have often focused on the Asian-lineage strains and have well-documented fetal growth dynamics (19, 51). However, in African-lineage infections, due to the increase in fetal demise, fetal growth dynamics have not been well characterized. Nevertheless, we measured growth dynamics and placental function to verify the extent of vaccination protection. Compared to historical controls from the Oregon National Primate Research Center (ONPRC,) all three endpoint cases (C3, V1, V3) had normal growth dynamics, including animal V1, which had a significant fluid collection **(Figure 7B-E)**.

**Figure 7.**
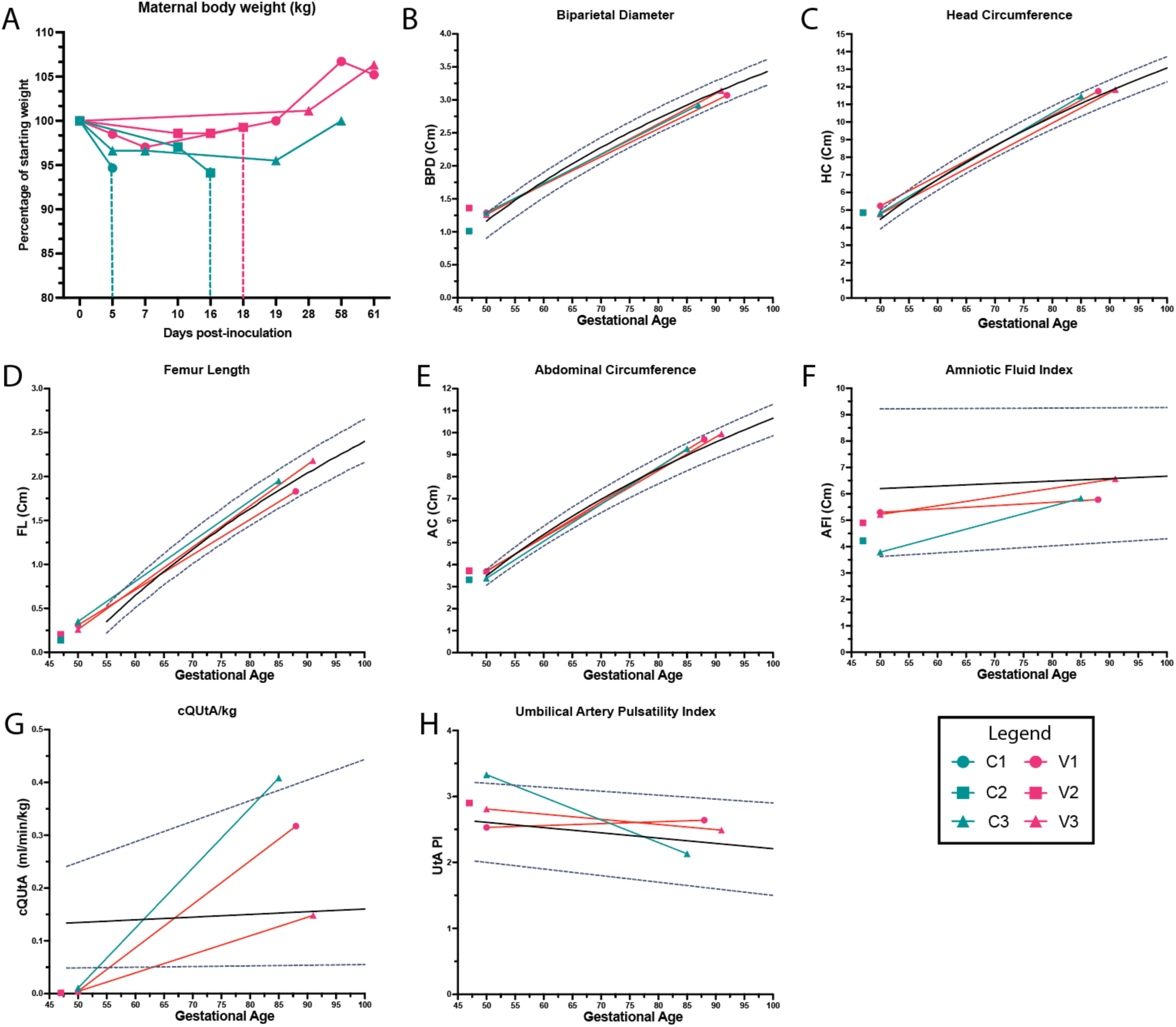
**Pregnancy dynamics measured for endpoint cases longitudinally show normal growth dynamics when compared to institutional controls. *A***, Maternal body weight was measured at indicated timepoints for each animal and graphed as a percentage of starting body weight. Longitudinal ultrasound measurements of fetal growth across gestation including *(**B**)* Biparietal Diameter (BPD), *(**C**)* Head Circumference (HC), *(**D**)* Abdominal Circumference (AC), and *(**E**)* Femur length (FL) in the six ZIKV-challenged fetuses with Alum control animals (n=3) in teal and VLP +Alum (n=3) in pink were plotted against historical controls, from ONPRC published data used to calculate the logarithmic regression for 50^th^ percentile (black solid line) and for 10^th^ and 90^th^ (dashed line) (52). ***F***, Amniotic Fluid Index data was obtained by standard measurement of four quadrants and plotted against historical control with the logarithmic regression representing for 50^th^ percentile (black solid line) and for 5^th^ and 95^th^ (dashed line). ***G***, UtA blood flow (cQtA/kg) corrected for maternal body weight using UtA diameter, maternal UtA cross-sectional area and volume of blood flow through the maternal UtA as previously described (43). ***H***, Doppler measurements were also used to calculate the umbilical artery Pulsatility Index (PI). Graphs were again plotted against ONPRC historical data. Graphs were created using GraphPad Prism v10.2.

To validate the fetal biometry data, we assessed amniotic fluid and uteroplacental hemodynamics for the placenta by Doppler ultrasound prior to infection and across gestation until 60 days post-inoculation (approximately GD90). Using the standard four-quadrant measurements to calculate amniotic fluid index, we observed no significant changes outside the range of historical controls for any of the end point cases **(Figure 7F)**. Corrected for maternal body weight, the calculated blood flow volume in the uterine artery (cQuta) demonstrated some variability between ZIKV animals but revealed an overall trend of appropriately increasing with advancing gestational age **(Figure 7G).**

Additionally, none of the umbilical artery pulsatility indices (PI) deviated from historical controls in any of the ZIKV+ animals **(Figure 7H)**. These data show that endpoint cases had normal pregnancy dynamics, suggesting that once animals make it past a certain gestational age, they are protected from demise, as seen in previous studies (19, 20).

### 2.6 vRNA tissue burden is highest among early demise cases

To assess systemic viral dissemination, we quantified ZIKV RNA (vRNA) in maternal tissues collected at necropsy. Animals that had clinical signs of early fetal demise, particularly those in the alum-only control group, had markedly elevated vRNA levels with up to 10^6^ across multiple organ systems. Up to 63 maternal tissues were tested at the time of necropsy for early demise cases (5-18 dpi) with detectable vRNA found in tissues from the lymph, musculoskeletal, digestive, endocrine, reproductive, central and peripheral nervous systems **(Figure 8; complete list of tissue samples in Table S2).** Overall, 38.09% of tissues from C1 and 30.16% from C2 were vRNA-positive compared to V2 which had 11.11% tissue positivity. Consistent with previous work, viral loads were highest in lymph tissues, particularly axillary and mesenteric lymph nodes (LN), that reached up to 1×10^6^ copies of vRNA per μg of RNA for all early demise cases **(Figure 8A)**. Control animals C1 and C2 had wide tissue distribution with genitourinary **(Figure 8B)**, musculoskeletal **(Figure 8C),** and gastrointestinal **(Figure 8D)** systems being positive with high levels of vRNA. In NHPs, ZIKV has been found in central nervous system tissue, although it is rare for adult human cases to develop encephalitis. Both central (basal ganglion) and peripheral (dorsal root ganglion) nervous tissue were positive in C1 and C2 though higher in C1 **(Figure 8E)**. Interestingly, dam V2 had detectable vRNA in LNs at lower levels with minimal tissue be due to previously acquired immunity. For pregnancies that did not end in demise, vRNA detection was minimal with C3 showing detectable vRNA in the jejunum, and V1 showing detectable vRNA in the fingers and wrist, which is a common finding in ZIKV infection. Viral RNA was not detected in tissues collected from V3 at necropsy which is consistent with the neutralizing activity seen prior to inoculation of this dam (Figure 5C). The timepoint at which tissues were assayed for vRNA between animals is an important distinction in this study. Early demise cases C1 and C2 still had detectable viremia whereas endpoint cases had completely resolved viremia 53 days prior to necropsy. Because this study was designed to compare fetal survival rather than viral loads between the vaccinated and unvaccinated cohorts, we cannot directly compare virus in tissues between survival and early demise cases. However, we note that maternal viremia was significantly lower in the animals that reached the study endpoint.

**Figure 8.**
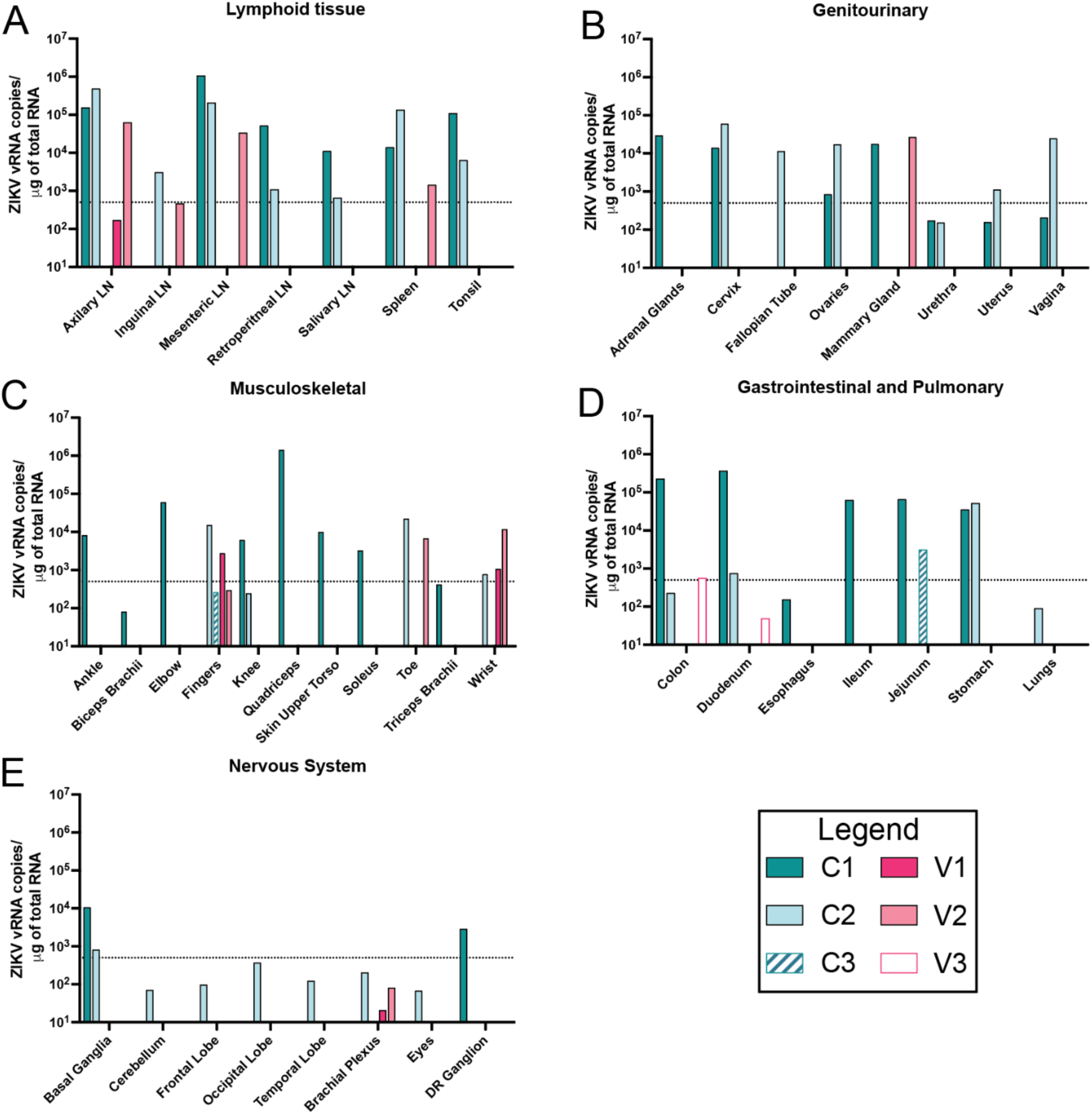
**ZIKV tissue distribution in dams. *A***, Graphical depictions of viral loads from **S. Table 2** were detected via one-step RT-qPCR. The limit of detection (LOD) was 500 copies of ZIKV vRNA. Graphs represent tissues that had at least one animal with positive viral detection and are separated into five tissue types. ***A-E***, Lymphoid, Genitourinary, Musculoskeletal, Gastrointestinal and Pulmonary, and Nervous System tissues were separated for each tissue. Graphs were created using GraphPad Prism v10.2.

**Table 2.** Placental viral loads. Total RNA was isolated from mapped placental cotyledons from both primary (1°) and secondary (2°) discs. All available cotyledons were tested; however, the number of cotyledons varied between placentas. ZIKV RNA levels were quantified using a one-step RT-qPCR assay in triplicate. Mean quantity values are based on standard ZIKV FFU and are presented as log10 viral copies per ml of tissue homogenate. The limit of detection was approximately 500 genomes (2.7). Positive values are indicated by a red gradient, values under the limit of detection are indicated in blue, no detectable value is denoted by —, and tissues not tested are indicated in gray.

|  | Alum-only |  |  |  |  |  | VLP + Alum |  |  |  |  |  |
| --- | --- | --- | --- | --- | --- | --- | --- | --- | --- | --- | --- | --- |
|  | C1 |  | C2 |  | C3 |  | V1 |  | V2 |  | V3 |  |
| Placental Disc | 1° | 2° | 1° | 2° | 1° | 2° | 1° | 2° | 1° | 2° | 1° | 2° |
| Cotyledon # |  |  |  |  |  |  |  |  |  |  |  |  |
| 1 | 3.67 |  | 7.68 |  | — | — | — | — | 1.28 |  | 2.79 | 1.45 |
| 2 | 5.74 |  | 7.59 |  | — | — | — | — | — |  | — | — |
| 3 | 3.86 |  | 7.31 |  | — | — | 3.50 | 3.08 | — |  | — | — |
| 4 | 3.05 |  | 7.59 |  | — | — | 3.38 | 4.02 | — |  | — | — |
| 5 |  |  |  |  | — | — | 3.26 | — |  |  | — | 4.56 |
| 6 |  |  |  |  | — | — | 3.22 | — |  |  | — |  |
| 7 |  |  |  |  | — | — | — | — |  |  | — |  |
| 8 |  |  |  |  | — | — |  | 3.81 |  |  | — |  |
| Decidua | 4.51 |  | 5.12 |  | — |  | — |  | 2.73 |  | 3.29 |  |
| Umbilical Cord | — |  | 6.66 |  | — |  | — |  |  |  | — |  |
| Amniotic Membrane | — |  |  |  | — |  | — |  |  |  | — |  |
| Chorionic Membrane | — |  |  |  | — |  | — |  |  |  | — |  |
| Total Membranes | — |  |  |  | — |  | — |  |  |  | — |  |

Although necropsies were performed at varying timepoints post-inoculation, prior studies have demonstrated that vRNA can persist at high levels (up to 1×10⁶ copies/μg RNA) as late as 60 dpi, suggesting that differences here could reflect true biological variance rather than timing alone (44). Among vaccinated animals that reached endpoint, vRNA burden was minimal (V1 6.5%) or no vRNA was detected (V3). This is consistent with earlier plasma viremia, which showed lower curves for the three animals that made it to endpoint, supporting the protective effect that ZIKV-VLP vaccination has against systemic viral dissemination.

### 2.7 Infectious viremia and placental damage is highest among the control group

To further investigate the relationship between maternal infection and fetal demise, we performed infectious virus isolation from select tissues on the day of necropsy. Maternal and placental tissues were homogenized, and the cell suspensions were co-cultured with *Aedes albopictus* C6/36 cells for five days. Focus forming assays were performed on supernatants collected from cocultures to quantify infectious virus, which was only detected in animals C1 and C2. C1 had high levels of virus detected in axillary LN cells and splenocytes whereas C2 had virus detected to a lesser extent. C2 additionally had high levels of infectious virus detected in villous placenta, decidua, and amniotic fluid **(Figure 9A)**. Although placental tissues from C1 were unable to be tested for infectious virus due to early loss at 5 dpi, vRNA levels in villous and decidual tissues were significantly elevated, suggesting the likely presence of replicating virus at the time of necropsy **(Table 2)**. Infectious virus was not detected in plasma samples at these timepoints, indicating that infectious virus was generated in the tissues.

**Figure 9.**
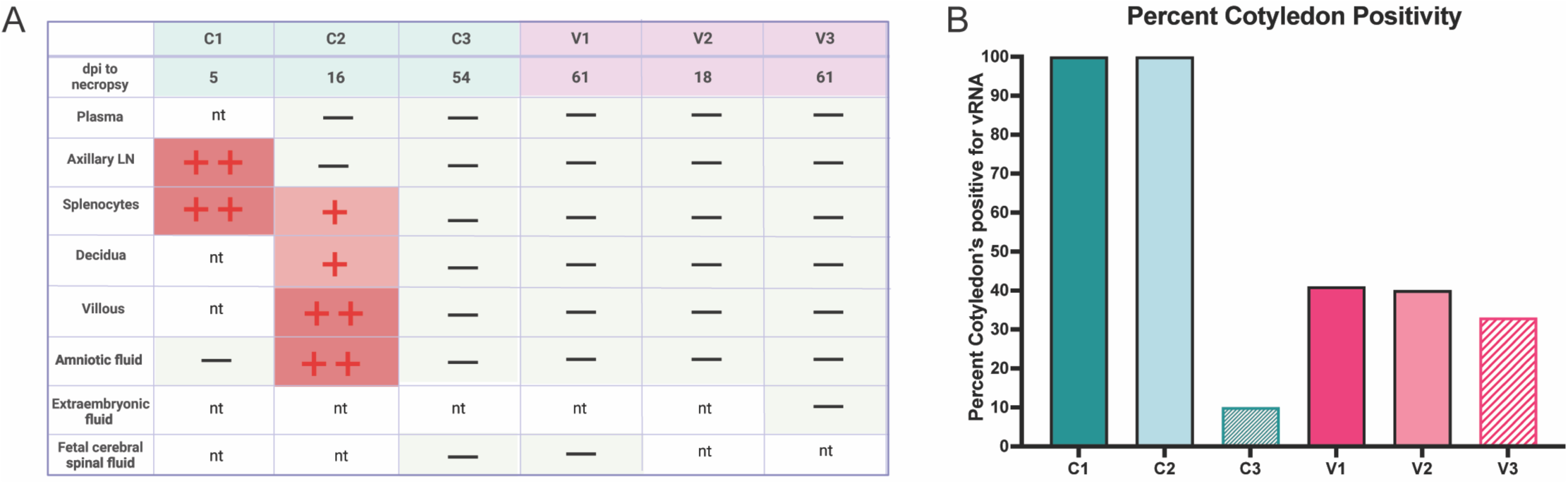
**Infectious virus and placental vRNA loads. *A,*** infectious virus was isolated from tissue homogenate, plasma, or fetal fluids. 400µl was added to C6/36 cells for 3 days and then titered on vero cells in a focus forming assay where + = virus detected (<10^5^ FFU/mL), ++ = high virus titer detected (10^5^-10^8^ FFU/mL), - = not detected, nt = not tested. Table created in BioRender. Jaeger, H. (2025) https://BioRender.com/4sig92p. ***B,*** Total RNA was isolated from mapped placental cotyledons and quantified using a one-step RT-qPCR for ZIKV RNA, summarized in Table 2. Percent positivity was determined by the number of cotyledons with vRNA detected over the LOD divided by total amount of cotyledons tested for each animal. Graph created using GraphPad prism v10.2.

Interestingly, we did not detect infectious virus or viral RNA from V2, which was an early demise case **(Figure 9A and Table 2)**. Similarly, no infectious virus was recovered from vaccinated or control animals that reached endpoint (V1, V3, and C3). This again is likely due to the timepoint of necropsy (∼60 dpi) where we do not expect to isolate infectious virus. Animal V1 had seven placental cotyledons with detectable vRNA, which associate in the absence of statistics with the dam plasma viremia observed **(Table 2)**. Animals C3 and V3 had lower positivity across the placenta, which is consistent with the lower or absent viremia seen in these two dams. To summarize the data, percent positive cotyledons were calculated for each animal that was above the limit of detection **(Figure 9B)**. These findings demonstrate that widespread infectious virus and elevated vRNA burden were restricted to unvaccinated animals with early fetal loss, supporting a link between uncontrolled maternal viral replication and placental infection.

### 2.8 Placental pathology reflects infection severity and fetal outcome

Placental infarctions, calcifications, decreased blood flow, and vasculitis are often found in ZIKV+ NHP models and human cases (42), which can be variable and, in some instances, have unknown effects on the long-term health of the infant. To investigate whether vaccination could protect against placental damage, histopathology was evaluated via blinded scoring of gross and histologic changes in individual cotyledons. Among early demise cases, control animals had the most prominent changes noted both on gross pathology and by histology. Gross evaluation of dam C1 revealed a marginal placental abruption affecting 25% of the primary placental disc **(Figure 10A, D)**, which was estimated to have occurred 1–2 days prior to necropsy. Histologically, this was associated with infarction and decreased oxygenation of the fetus, evident by widespread elevation of nucleated fetal red blood cells (1–2 per high-power field, **Figure 10G**), an abnormal finding at this gestational stage. In contrast, control dam C2 and vaccinated dam V2, both of which developed hydrops fetalis, had minimal gross placental abnormalities but showed histologic signs of infection and hypoxia. C2 had infarctions on gross examination, indicated by the white asterisks **(Figure 10B)**. Microscopically, both animals (C2, V2) had elevated nucleated fetal red blood cells (2-3 per high-powered field) with nuclear clearing **(Figure 10H)**, a feature associated with parvovirus B-19 infection. However, all three early demise animals tested negative for parvovirus B-19 via antibody staining **(S Figure 7)**, suggesting that the nuclear clearing was more likely due to hypoxia or ZIKV-induced placental dysfunction which can also cause this histological feature. Additional findings in C2 included calcifications and leukocytoclastic vasculitis in two maternal spiral arteries indicated by the black asterisks **(Figure 10E)**, and minimal apoptotic debris **(S Figure 8A)** further indicating significant placental inflammation and vascular compromise within this animal.

**Figure 10.**
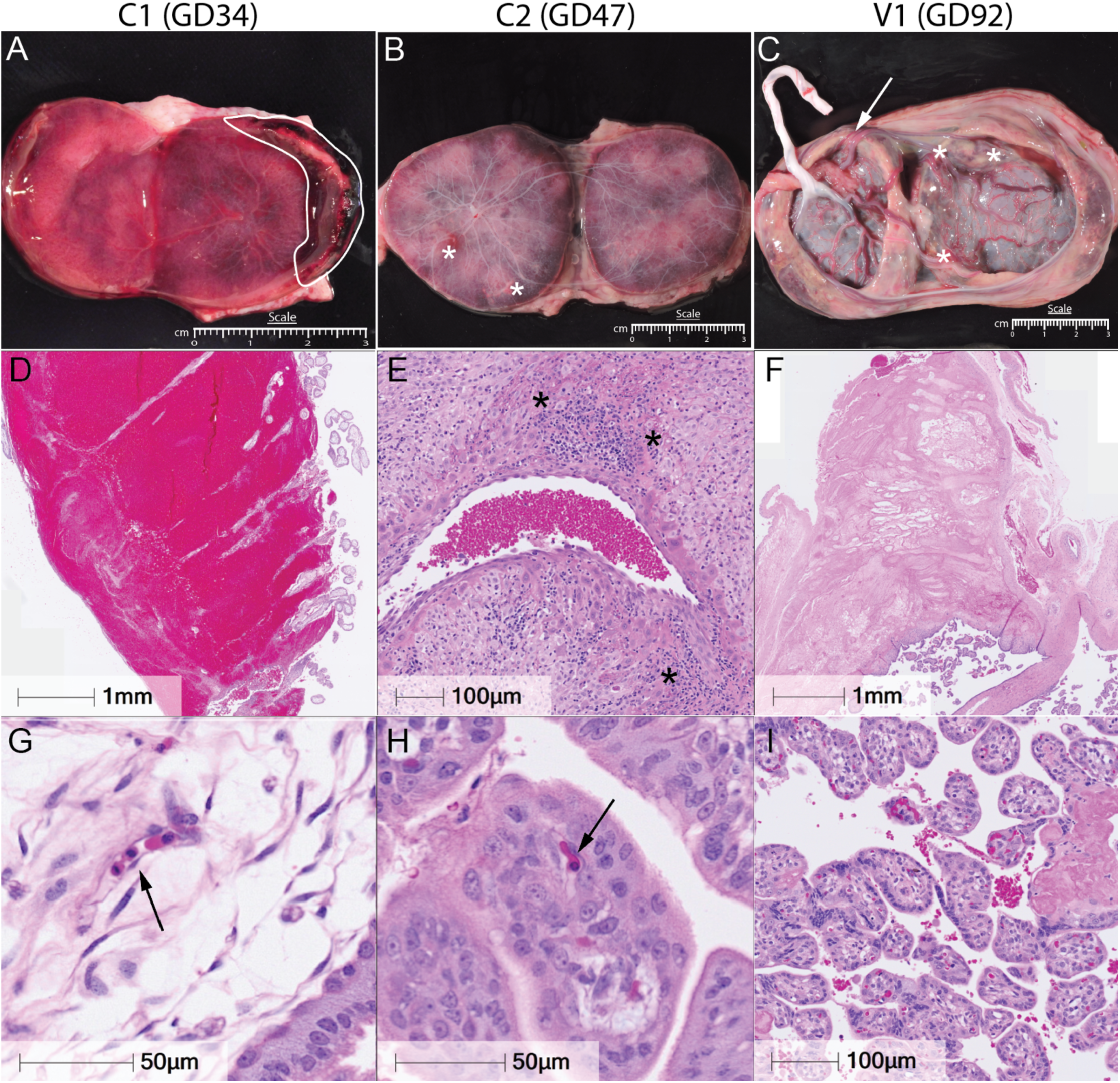
**ZIKV infection induces placental abnormalities including infarction, abruption, and vascular lesions in early demise cases**. Gross and histopathological examination of placentas from animals C1 *(A, D, G)*, C2 *(B, E, H)*, and V1 *(C, F, I)*, collected at necropsy on GD34, GD47, and GD92, respectively. C1 *(A)* shows a large placental abruption outlined in white, consistent with an event occurring within 24 hours prior to delivery. In C2 and V1 *(B, C)*, white asterisks mark infarctions, and a white arrowhead in V1 *(C)* indicates a circumvallate placenta. The corresponding hematoxylin and eosin-stained sections are shown in *D–I,* where the abruption identified grossly in C1 is confirmed histologically *(D)*, C2 shows leukocytoclastic vasculitis and spiral artery remodeling (E, black asterisks), and V1 demonstrates a microinfarction *(F)*. Additional histopathological features are shown in *G–I,* where C1 contains red blood cells (RBCs) without nuclear clearing indicated by the black arrowhead *(G)*, C2 displays RBCs with nuclear clearing *(H)*, black arrowhead), typically observed at 2–3 per high-power field, and V1 exhibits moderate villous knotting *(I)*. Scale bars are shown in black; all histology images were acquired at 40× magnification.

The endpoint cases had relatively minimal pathology in comparison to early demise cases. Dam V1 showed signs of circumvallate placenta, separated membranes, and focal infarctions on ultrasound and gross pathology **(Figure 10C)**, with diffuse microinfarctions and sporadic villous knotting (**Figure 10F,I)**, likely occurring weeks prior to necropsy. These findings have a lower incidence of infarction and placental damage than our previous reports from GD90 ZIKV-infected placenta (42). Placentas from C3 and V3 were largely unremarkable, with only minor calcifications and focal microinfarctions noted in V3, a finding also seen in normal pregnancies **(S Figure 8B,C)**. Together, these data indicate that severe placental pathology and infectious virus recovery were limited to unvaccinated animals with early fetal demise. Vaccinated animals had either minimal or undetectable pathology and no infectious virus, supporting the protective impact of adjuvanted ZIKV-VLP vaccination against placental infection and associated damage.

### 2.9 Evaluation of ZIKV-DAK burden in fetuses

To investigate potential causes of fetal demise, we performed in situ hybridization (ISH) to assess ZIKV RNA distribution in whole-fetus sections from early demise cases, selected brain regions from endpoint cases, and the placenta. For cases of early demise, whole fixed fetuses were sagittally bisected along the midline prior to paraffin embedding. Sections were collected from both sides of the fetus, and one “screening” slide was initially evaluated to determine which side contained the most representative tissue architecture for ISH analysis. In endpoint cases (i.e fetuses delivered at GD 90), a similar screening approach was applied to brain tissue to identify regions most likely to harbor viral RNA. Based on this screening, three brain regions—the frontal lobe, deep brain structures (including the basal ganglia), and cerebellum—were selected for targeted ISH staining.

For the endpoint cases we were able to detect vRNA via RT-qPCR on select tissues, excluding brain tissue **(S Table 3)**. Only the adrenal glands and submandibular region of C3F were positive for ZIKV RNA and the two vaccinated animals were completely negative. At this timepoint post-inoculation it is common to see 5-8 tissues positive for ZIKV (42, 43, 45). For endpoint brain tissue, ISH was used, and screening slides revealed almost no detectable ZIKV RNA **(S Figure 9A)**. To confirm these findings, each selected brain region underwent further sectioning, with four additional slides cut at 10 μm intervals from the screening plane and processed for ISH. This approach verified that vRNA was largely absent in these tissues. In V1F, a small number of apparent positive cells were observed, but their morphology and distribution suggested they were staining artifacts rather than true viral signal **(S Figure 9B)**. A similar result was seen in early demise cases C1F and V2F **(S Figure 9C,D)**.

Minimal ZIKV RNA detection was observed in fetus C1F fetal and placental tissues, consistent with the early timing of demise (day 5 post-infection) **(S Figure 9C)**. This aligns with prior studies showing that ZIKV typically requires 7-10 days post-infection to cross the placental barrier and establish productive fetal infection, suggesting that the widespread viral dissemination observed in C1F was in line with the natural infection progression (23).

Interestingly, no ZIKV RNA was detected in the vaccinated early demise fetus V2F or its placenta, which demised due to hydrops fetalis as well. This finding suggests that either a non-viral process contributed to the fetal loss or that infection had not yet established within the fetus, and instead, systemic or placental pathology (e.g., hypoxia) may have been the proximate cause. In contrast, fetus C2F exhibited clear ZIKV RNA positivity across multiple tissues **(Figure 11A-C)**. This finding revealed widespread infection, with viral loads detected in the lungs, heart, kidney, gastrointestinal tract, liver, musculoskeletal tissues, and multiple brain regions, including the frontal lobe, meninges, and soft tissues surrounding the skull **(Figure 11D-L)**. Halo AI-assisted image analysis was employed to quantify the proportion of infected cells within each organ, providing an objective and reproducible measure of viral burden. For quantification, five serial slides were developed at 5 μm intervals, with each data point on the graph representing an individual slide **(Figure 11M)**.

**Figure 11.**
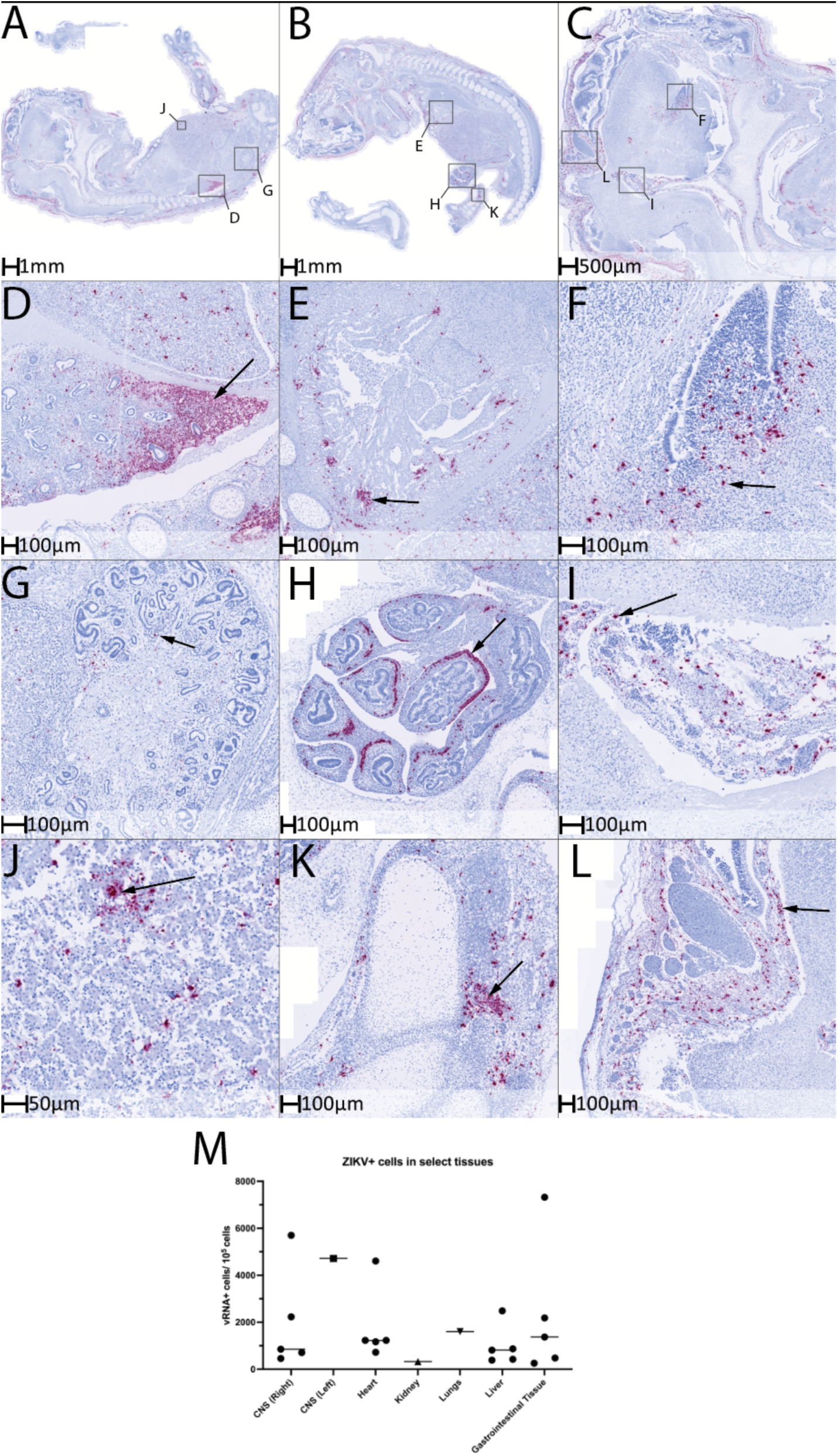
**ZIKV vRNA was detected via ISH staining only in case C2. *(A–C)*** Representative sagittal sections of FFPE-blocked C2 fetus stained with a ZIKV-specific ISH probe (Warp Red) and counterstained with hematoxylin and lithium blue. More intact brain regions were visible on the left side ***(A, C)***, while major organs and musculoskeletal structures were better preserved on the right. Each column corresponds to the low-magnification overview images shown in ***A–C***. ZIKV+ cells were detected in multiple major organs, including the lungs ***(D)***, heart ***(E)***, kidney ***(G)***, gastrointestinal tract ***(H)***, liver ***(J)***, and musculoskeletal regions such as the foot ***(K)***. Viral RNA was also detected throughout the central nervous system, including the frontal lobe ***(F)***, meninges **(I)**, and skin and soft tissue surrounding the skull. Scale bars are shown in black; all histology images were acquired at 40× magnification. ***(M)*** Quantification of total cell counts (blue nuclei) and ZIKV+ cells (Warp Red staining) was performed using Halo AI (Indica Labs).5 sections were stain from right side (B) where the 5 data points represent 5 separate slides. The kidney, lungs and brain were quantified from the left side and only have one data point representing the one slide that was stained. Graph was created using GraphPad Prism v10.2.

### 2.10 Evidence of hypoxia and cardiovascular compromise in hydrops fetalis Cases

To determine the etiology of fetal hydrops and demise in our study, we conducted histological and immunohistochemical analyses on fetal and placental tissues from hydrops cases (C2F and V2F), focusing on markers of fibrosis, angiogenesis, and immune infiltration. Serial 5 μm sections were stained with Trichrome, CD31 (endothelial marker), and CD68/CD163 (macrophage markers) to assess tissue structure, vascular injury, and immune cell localization **(Figure 12)**.

**Figure 12.**
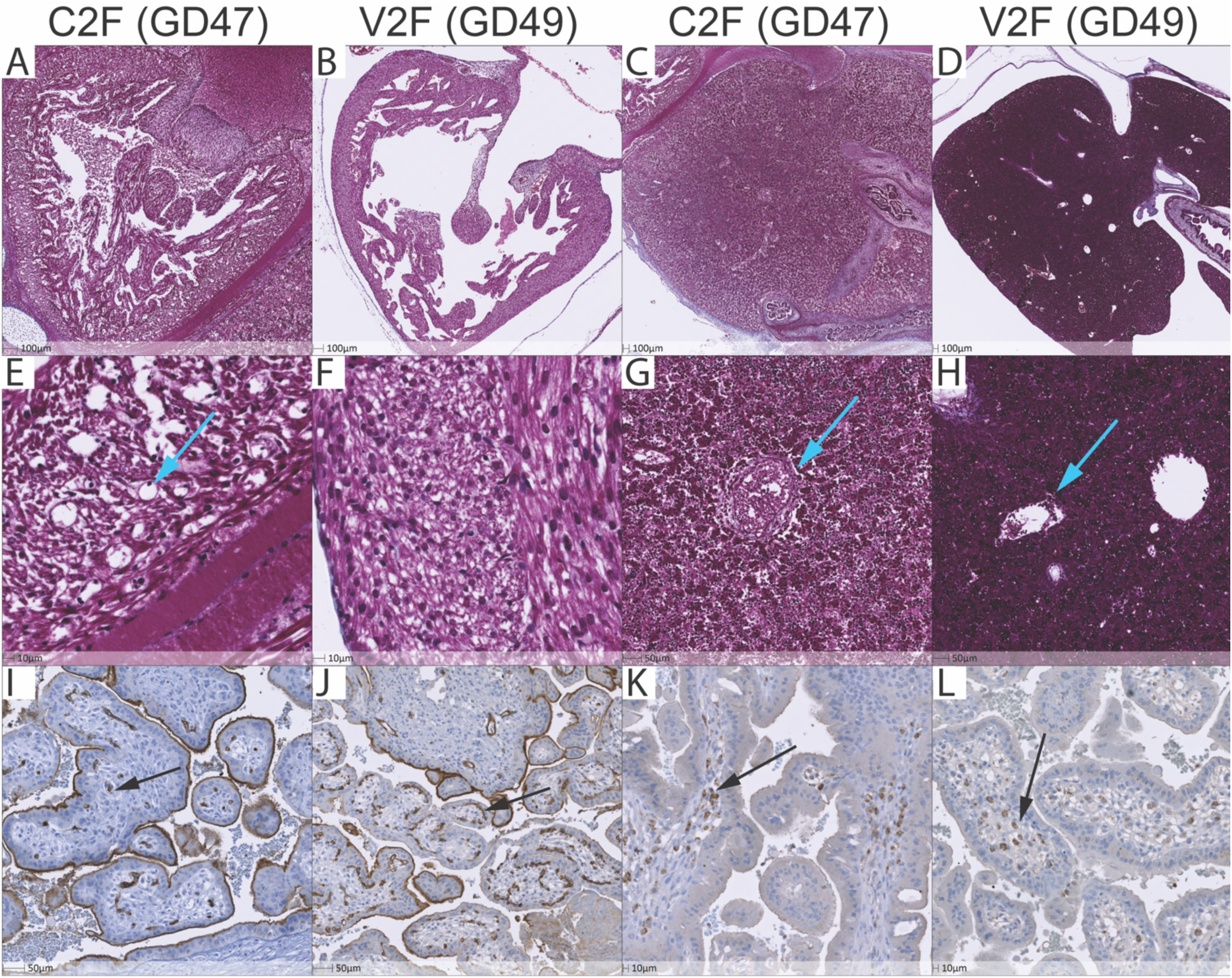
**Hydrops fetalis cases show signs of hypoxia. *(A–H)*** Representative sagittal sections from formalin-fixed paraffin-embedded (FFPE) fetuses C2F **(A, C, E, G)** and V2F **(B, D, F, H)** stained with Masson’s Trichrome. **(A, B)** Low-magnification images of the heart; corresponding high-magnification views shown in **(E, F)**. Cyan arrow in **(E)** indicates a proangiogenic response within the C2 fetal myocardium. **(C, D)** Low-magnification images of the liver; corresponding high-magnification views shown in **(G, H)**. Cyan arrows in **(G, H)** identify portal vessels lacking fibrosis—absence of blue staining confirms minimal collagen deposition. **(I, J)** Fetal placental sections stained for CD31 with DAB chromogen (brown). Black arrows indicate fetal capillaries within the villous stroma. **(K, L)** Fetal placental sections stained for CD68 with DAB chromogen (brown). Black arrows highlight fetal macrophages (Hofbauer cells) within the villous stroma. All images acquired at 40× magnification; scale bars shown in black.

Although the pathogenesis of hydrops fetalis remains incompletely understood, it is broadly linked to multifactorial disturbances in fetal or placental physiology. Both low- and high-output cardiac failure can cause elevated central venous pressure, which in turn increases capillary hydrostatic pressure and impairs lymphatic return to the vascular space (53, 54). In C2F, where viral RNA was detected extensively, including within the myocardium (**Figure 11E**), we hypothesized that direct viral injury could have led to low-output heart failure. However, Trichrome staining of the heart and liver did not reveal significant fibrosis around the portal triad for both animals **(cyan arrows),** suggesting a more acute or inflammatory process rather than chronic heart failure **(Figure 12C, D, G, H)**. Trichrome staining did reveal a proangiogenic response within the myocardium, consistent with compensatory remodeling in response to hypoxic stress that was seen extensively in C2F, but not in V2F **(Figure 12**

To further explore our hypothesis of hypoxia-driven vascular injury we stained serial placental sections with ZIKV-ISH, CD31 and CD68/163. Hypoxia can induce endothelial injury and capillary dropout resulting in increased vascular permeability and interstitial fluid accumulation (55). Again, V2F showed no ZIKV positivity via ISH with extensive staining seen in C2F fetal placenta **(S Figure 10A-D).** Despite lacking detectable vRNA, V2F had clear histological evidence of placental hypoxia, including loss of villous stromal capillaries and endothelial injury confirmed by CD31 staining **(Black arrows, Figure 12J)** when compared with C2F that had relatively normal fetal capillary distribution **(Figure 12I)**. Additionally, CD68+ Hofbauer cells were present at increased density in both C2F and V2F, suggesting an inflammatory component, although not to the same extent as seen with chronic ZIKV infection in Asian-lineages **(Figure 12K,L)**. Histologically, elevated nucleated fetal red blood cells and placental vascular injury further supported a hypoxic pathology for both C2F and V2F. Together, these findings demonstrate histological evidence of placental hypoxia and vascular injury in both C2F and V2F, independent of detectable vRNA.

## 3. Discussion

In this study, we evaluated the immunogenicity and protective efficacy of a ZIKV-VLP vaccine adjuvanted with Alhydrogel (alum) in mice, nonpregnant and pregnant rhesus macaques. Our findings demonstrate that the ZIKV-VLP vaccine elicited robust humoral responses in both mice and nonpregnant rhesus macaques, additionally limiting viral dissemination in macaques. In pregnant rhesus macaques, the VLP conferred partial protection from early demise and viral dissemination. The ZIKV-VLP vaccine builds on prior successes of flavivirus VLP-based platforms, which have demonstrated potent induction of neutralizing antibodies and protective efficacy in preclinical models of arboviruses (39, 41, 56). The alum-adjuvanted formulation used here generated neutralizing antibody titers in both mice and rhesus macaques. In pregnancy, while vaccinated dams experienced reduced viremia, lower placental and fetal viral burden, and improved fetal survival compared to controls, fetal loss still occurred in one case. We note that neutralizing antibody titers had fallen between the time of final boost and inoculation, suggesting that using other adjuvant systems or increased amounts of vaccine antigen might promote greater vaccine efficacy, as we have seen with other VLP-based arboviral vaccines. These results underscore the unique immunologic challenges of pregnancy and highlight the need for vaccine strategies specifically tailored to this setting.

### Use of African-lineage ZIKV-Dakar

Several recent studies have modeled a frequent preterm pregnancy loss at a rate of 78% following challenge with the African-lineage strain Dakar-41524 (ZIKV-DAK) (21, 23, 57). This rate is substantially higher than the background incidence of 4–10% fetal loss typically observed in rhesus macaque colonies, as well as the 16.7% loss reported across primate centers following infection with Asian-lineage strains at gestational day 30 (n=9), where the overall fetal loss rate across ZIKV strains and gestational timepoints was 26% (20). The mechanism of ZIKV-induced pregnancy loss is not fully characterized as the number of documented ZIKV-induced fetal demise cases in humans is low (46, 47). The use of the African-lineage ZIKV-DAK provides new insights into ZIKV-induced fetal loss and provides a higher penetrance model for the evaluation of vaccines and prevention strategies. Previous NHP studies using the African strain ZIKV-DAK point to ZIKV gaining access to the fetal compartment through paraplacental transmission between 7-9 dpi (23). This event happens through infection at the fetal membranes and subsequent dissemination to the villous stroma that occurs around day 6 or 7 (23). In our study, we observed a similar rate of fetal loss in our control cohort of 66% with fetal loss occurring at 5-16 dpi. Similar to previous studies, we found extensive ZIKV RNA via ISH in the placental stroma in C2F (14-16dpi) and none in C1F (5dpi). These findings reinforce the utility of ZIKV-DAK as a stringent, high-penetrance model for evaluating candidate vaccines and advancing our understanding of ZIKV-induced pathogenesis.

### Insights into mechanisms underlying hydrops fetalis in ZIKV-inoculate fetuses

Two fetal demise cases developed early hydrops fetalis. Both C2F and V2F exhibited ultrasound findings consistent with hydrops fetalis, including scalp edema, pleural effusion, and ascites at 14-16 dpi and 16–18 dpi, respectively. In C2F, which experienced in utero demise, extensive viral RNA was found within a range of major organ systems, and histological analyses revealed hallmark features of placental hypoxia. In contrast, V2F, which was electively necropsied between 16–18 dpi, no ZIKV RNA was detected, yet some histological features of hypoxia were observed. While these findings may indicate a non-viral component of fetal demise for V2F, dam V2 had higher peak viremia than animals that reached endpoint. Although fetal hydrops has been reported in ZIKV-infected human pregnancies and rarely in NHP ZIKV studies, the underlying mechanisms remain poorly understood (46, 47). Our study provides new insights into the pathogenesis of ZIKV-induced hydrops, implicating placental dysfunction and fetal hypoxia as central drivers. Collectively, our findings suggest that hydrops fetalis in ZIKV-infected pregnancies may arise from progressive hypoxic injury originating at the maternal-fetal interface, rather than solely from direct viral invasion.

To contextualize these findings, hydrops fetalis in humans is described as either immune or nonimmune hydrops fetalis (NIHF). Common causes of NIHF include chromosomal abnormalities, infections, placental pathology, and cardiovascular issues, with chromosomal and, less frequently, cardiovascular anomalies being the most likely etiology in the first trimester (58). Heart failure can also be caused by Parvovirus B19 infection via severe fetal anemia and sometimes direct myocarditis leading to high-output cardiac failure. This, in turn, causes fluid accumulation within the interstitial space. Although anemia is a key feature in parvovirus-driven hydrops, heart failure is often the terminal mechanism, precipitated either by demand or direct myocardial injury. Similarly in our study, pregnancies complicated by hydrops fetalis also displayed several other effects including severe edema, hepatomegaly, and placental vasculopathy. However, we did not detect evidence of viral-induced anemia nor Parvo-B19 antibodies on immunohistochemistry. Instead, histology showed prominent placental infarctions, vasculitis, and nucleated fetal red blood cells, consistent with hypoxia due to placental insufficiency.

### Correlates of protection

In our study, vaccination reduced maternal viremia and mitigated the severity of placental pathology in surviving pregnancies but did not fully prevent the development of hydrops fetalis or fetal demise. The timing of challenge post-prime in the pregnancy cohort was substantially longer than the nonpregnant cohort which could contribute to lower neutralizing responses pre-challenge. This suggests that additional immune mechanisms beyond systemic humoral responses may be required to achieve sterilizing protection at the maternal–fetal interface. Local immunity within the placenta, including the activity of tissue-resident immune cells and the efficiency of transplacental antibody transfer, may play a critical role in limiting viral access to the fetus. Furthermore, the timing of immunization relative to gestation could also influence efficacy, given the defined window of placental susceptibility to viral invasion observed with ZIKV-DAK. Importantly, we did not observe an association between vaccination-to-challenge interval and fetal outcome, as the two vaccinated dams that reached endpoint were challenged at 206- and 154-days post-vaccination, whereas the early demise case (V2F) was challenged at 198 days. Immune parameters such as Fc-mediated effector functions, mucosal immunity within the reproductive tract, or induction of placental antiviral pathways may also contribute to protection and warrant further investigation. Given the absence of ongoing large-scale ZIKV outbreaks, the use of high-stringency NHP models such as ZIKV-DAK will remain essential for defining correlates of protection and guiding the development of vaccine platforms capable of providing comprehensive maternal and fetal protection.

## 4. Conclusions

The results from this study suggest important considerations for immunization strategies that provide comprehensive protection during pregnancy. These conclusions highlight the need to fully assess the efficacy of vaccine candidate to ensure full protection against early pregnancy loss secondary to ZIKV infection. Our findings suggest that, while vaccination can protect against viremia in a nonpregnant model, these responses do not translate to complete protection from fetal demise during pregnancy. While the VLP-based vaccine approach described here required further optimization (*e.g.* a more effective adjuvant), balance of a safe vaccine platform that provides durable protection is a concept that should be carefully considered when developing ZIKV vaccines.

## 6. Methods

### 6.1 Animal ethics statement

All experimental protocols were approved by the Institutional Animal Care and Use Committee (IACUC) of the Oregon National Primate Research Center (43), which is accredited by the Assessment and Accreditation of Laboratory Animal Care (AAALAC) International. All procedures in this study complied with the ethical standards outlined in the Animal Welfare Act, as enforced by the United States Department of Agriculture, and followed national guidelines for the care and use of laboratory animals. The study also adheres to the ARRIVE guidelines for animal research (59).

### 6.2 Zika virus (ZIKV) preparation and cell culture

Zika virus (ZIKV) strain PRVABC59 (ZIKV-PR) was provided by the Centers for Disease Control (CDC) and prepared as previously stated (42–45). Briefly, the virus was passed twice on C6/36 cells (ATCC CRL-1660), and a working stock was concentrated by ultracentrifugation through a 20% sorbitol cushion and titered in Vero cells (ATCC CRL-1586). ZIKV-DAK used in the pregnancy challenge study was provided by Dr. Thomas Friedrich at the University of Wisconsin as previously described (21–23, 57, 60–62). Both working stocks were sequenced prior to use. All cells were cultured in Dulbecco’s modified Eagle medium (DMEM) containing 2 mM L-glutamine (Invitrogen, Waltham, MA), 100 U/ml penicillin G-sodium, 100 μg/ml streptomycin sulfate (Invitrogen, Waltham, MA), and 10% fetal calf serum (FCS).

*6.3 Virus- like particle (VLP) production*

The prM-Eregion of ZIKV PRVABC59 (GenBank accession MH158237.1) was expressed either from a replication-deficient adenovirus or from a stable cell line that inducibly expresses prM-E. For the adenovirus, the ZIKV genome encoding anchor region of C through the end of E (nucleotides 396 to 2489) was amplified by RT-PCR and cloned into pAdtet7 (PMID: 10470081) downstream of the tetracycline (tet) operator. The resultant vector was transfected into 293-cre cells that were subsequently infected with an E1– adenovirus helper virus (Ψ5) that has loxP sites flanking the DNA packaging signal. A recombinant virus, AdtetZprM-E, was derived from cre-mediated recombination between the shuttle plasmid and the helper virus. AdtetZprM-E and a second adenovirus, Adtet-trans, that expresses the tet transactivator protein, were used to infect Vero cells. ZIKV-VLP were collected from culture supernatants at 3-4 d pi. For the cell line, the prM-E region of ZIKV (nucleotides 468-2489) was cloned in frame behind the tissue plasminogen activator signal sequence (PMID: 33762420) into vector pLVX-Tight-Puro (Takara Bio). This vector was stably transfected into HEK293 cells that constitutively express the tet-ON transactivator. Puromycin resistant cell lines were treated with 1 µg/ml doxycycline (dox) and analyzed for E expression. For VLP production, cells were treated with dox and VLPs collected from culture supernatants at 5-7 days post dox addition. For both adenovirus and cell lines, supernatants were passed through 0.45 µM filters and further clarified by low speed centrifugation. VLPs were then concentrated and purified by ultra-centrifugation through a 20% sorbitol 50 mM Tris 1 mM MgCl_2_ pH 8.0 buffer at 150,000 x g for 2 hours at 4°C. Pellets were then resuspended in 1/100th of the original volume with 10 mM Tris 150 mM NaCl buffer supplemented with 5% trehalose for protein stability and stored at −80°C until use.

### 6.4 Western blot

Detection of viral protein was performed by immunoblotting with flavivirus envelope-specific monoclonal mouse anti-4G2 (ATCC: HB-112) to visualize the E-protein. Quantitation of envelope protein on immunoblots was determined using titered ZIKV-PR stock to determine focus forming unit equivalents per mL.

### 6.5 Mouse experiment

C57BL/6J mice (Jackson Laboratories) were injected intramuscularly with a prime-boost method (D0 and D21) with 10^6^ focus forming equivalents of VLP alone or adjuvanted with 150 μg Alhydrogel (Invitrogen vac-alu-50). Serum was collected at 6 weeks post-vaccination.

### 6.6 Non-pregnant vaccine challenge experiment

Rhesus macaques (*Macaca Mulatta*) (n = 14) used in this safety, immunogenicity, and efficacy study were healthy, average-weight animals that were fed a standard chow diet (Purina Mills) and had access to water ad libitum. Animals were randomly assigned PBS control (n=3; 16yr. female, 6yr male, 6yr male), VLP alone (n=6, 13yr male, 14yr female, 6yr male, 5 yr male, 10 yr female, 16yr female), or VLP + Alum (n=5; 6yr male, 12yr female, 6yr male, 7yr male, 10yr female, 8yr female) group with age- matched and sex distributed animals in each group to control for age and sex as a variable. RM were vaccinated subcutaneously (s.q.) using a prime-boost method, receiving a boost 21 days apart. Blood, urine, and clinical assessments were performed on days 0, 1, 2, 3, 4, 5, 7, 11, 15, 22, 28 dpi. All animals were monitored closely for changes in behavior, appetite, and weight. Animals were humanely euthanized at 28 dpi and select tissues were taken to monitor viral dissemination.

### 6.7. ZIKV enzyme-linked immunosorbent assay (ELISA)

Enzyme-linked immunosorbent assays (ELISAs) were conducted on high-binding 96-well plates (Corning) coated for 3 days at 4°C with 1×10⁶ FFU per well of purified ZIKV-PR in PBS. After blocking with 5% milk in PBS containing 0.05% Tween-20, serial five-fold dilutions of heat-inactivated plasma were added, starting at a 1:50 dilution. Detection was performed using horseradish peroxidase (HRP)- conjugated anti-monkey IgG or IgM secondary antibodies (Rockland) at 1:1000, followed by OPD substrate (0.05 M citrate buffer, 0.4 mg/mL o-phenylenediamine, 0.01% hydrogen peroxide, pH 5; Life Technologies). After 5–7 minutes, the reactions were stopped with 1 M HCl. Absorbance was measured at 490 nm using a Synergy HTX Microplate Reader (BioTek, Winooski, VT) within 10 minutes of substrate addition. Endpoint titers were calculated using a Log/Log transformation method, and results were analyzed and graphed in GraphPad Prism v10.2 software.

### 6.8. Focus reduction neutralization (FRNT) assays

Focus Reduction Neutralization Tests (FRNTs) were conducted as previously described (42) to assess plasma antibody neutralizing activity in both nonpregnant and pregnant cohort studies. Heat- inactivated plasma samples were serially diluted five-fold in 2% FBS/DMEM, starting at a 1:20 dilution, and incubated with 200 FFU per well of ZIKV-PR at 37°C for 2 hours. The virus–plasma mixtures were then transferred onto monolayers of Vero cells in 96-well plates and incubated for 1 hour with rocking. After infection, cells were overlaid with carboxymethylcellulose (CMC) diluted in 2% FBS/DMEM and incubated for 30 hours. Plates were then fixed with 4% paraformaldehyde and blocked with 0.05% Tween-20 in PBS for 1 hour. To visualize ZIKV-infected foci, plates were stained with the flavivirus envelope-specific monoclonal antibody 4G2 (ATCC: HB-112), followed by HRP-conjugated anti-mouse IgG (Santa Cruz Biotechnology, sc2005). Foci were developed using VIP peroxidase substrate (Vector Laboratories) and quantified with an ELISpot reader (AID). The 50% focus reduction neutralization titers (FRNT₅₀) were calculated by non-linear regression in excel and graphed using GraphPad Prism v10.2, based on the percentage of foci remaining at each dilution relative to virus-only control wells.

### 6.9. ZIKV RNA detection

RNA was isolated from plasma, urine, and cerebrospinal fluid (CSF) using Promega Maxwell RSC viral TNA (Madison, Wisconsin) according to the manufacturer’s instructions. RNA from tissue samples were isolated using Zymo Research Direct-zol RNA Miniprep Kit (Cat. R2050) according to the manufacturer’s protocol. Tissue sections were collected into tubes containing 1 mL of TRIzol and approximately 250 μL of silica beads (VWR 48300–437). The samples were homogenized using a Precellys 24 homogenizer (Bertin Technologies) pulsing for two cycles of 45 seconds on and 30 seconds off. ZIKV RNA levels were measured by a one-step quantitative real-time reverse transcription polymerase chain reaction assay (RT-qPCR) using TaqMan fast virus One-Step RT-PCR Master Mix (Applied Biosystems) (44). Primer/probe sequences for the detection of ZIKV-PR (nonpregnant cohort) include Forward: 5’-TGCTCCCACCACTTCAACAA; Reverse: 5’TGA GGCAGGGAACCACAATGG; and TaqMan probe: 5’ Fam- TCCATCTCAAGGACGG -MGB(43). Primer/probe sequences for the detection of ZIKV-DAK (Pregnant cohort) include Forward: 5’- AARTACACATACCARAACAAAGTGT; Reverse: 5’ TCCRCTCCCYCTYTGGTCTTG; and TaqMan probe: 5’ FAM-CTYAGACCAGCTGAAR-MGB(63). For RNA standards, RNA was isolated from purified, tittered stock of ZIKV-PR or ZIKV-DAK.

### 6.10. Pregnant dam vaccination and time-mated breeding

Female rhesus macaques (n = 10) used in this study were all healthy, average-weight animals that were fed a standard chow diet (Purina Mills) and had access to water ad libitum (S. Table 1). The cohort size was determined based on our initial Zika virus studies by our group using similar assessments with the experimental design shown in **Figure 2** (42, 45). Animals were randomly assigned into Vaccination or Alum control groups with age matched animals in each group. Female animals were vaccinated using a prime-boost method, receiving a boost 21 days apart. All dams received a secondary booster to match the nonpregnant cohort IgG levels indicated in **Figure 1**. After vaccination or sham vaccination, dams were assigned to the ONPRC time-mated breeding (TMB) program. Daily blood sampling was used to coordinate the breeding attempts at 7 days prior to estradiol surge (42). Animals were then housed in female pairs with continuous full contact, except for the time (typically 3-5 days) that females were transferred and co-housed with the male breeder for TMB. All pregnancies were identified early by daily hormone monitoring and confirmed by prenatal ultrasound. The timed mating scheme allowed for accurate age determination within 24-48 h of conception. Four animals did not conceive after multiple breeding attempts and were withdrawn prior to challenge; resources were reallocated to enable deeper phenotyping of the remaining six animals (final n=6).

After pregnancy confirmation at gestational day (GD) 30, vaccinated dams were inoculated subcutaneously with a total of 1×10^5^ FFU of ZIKV-DAK diluted in 1ml of physiologic saline. The inoculum was distributed as ten total 100 µl injections at the radial and medial aspects of the wrist, the dorsal hand in both radial and medial positions, and one injection near the proximal antebrachium of each arm. This anatomical distribution was selected to promote uniform subcutaneous absorption and systemic dissemination of the virus, while ensuring consistency across all experimental groups.

### 6.11. Post-challenge clinical monitoring and endpoints

The animals were monitored daily for clinical signs of disease or discomfort in addition to cage- side ultrasound checks to monitor fetal cardiac activity. As depicted in Figure 2, ultrasound studies were performed to monitor fetal development on 21 and 60 dpi. Clinical assessments and maternal blood and urine collections were performed on days 0, 1, 2, 3, 5, 7, 10, 14, 21, and 28 dpi. To minimize sedation times, all fetal cardiac monitoring ultrasound checks were conducted without sedation or coordinated with blood draws when necessary to minimize stress by board-certified Maternal-fetal Medicine physician (J.O.L.) and a board-certified laboratory animal specialist veterinarian (J.S.). These procedures were conducted in accordance with established protocols at our institution to minimize stress and adverse outcomes (42, 52). Post-infection they were single-housed but given visual contact to prevent confounding by re-infection. All animals were monitored closely for changes in behavior, appetite, and weight. No signs of maternal distress or fetal loss were observed that could be attributed to the anesthesia or fasting regimen.

Our established protocols in maternal and fetal health were designed to determine clinical endpoints for humane euthanize. Three animals met this criterion due to fetal parameters. C1 showed partial cervical dilation on US, uterine contractions on exam, fetal bradycardia, and a hyperechoic placenta which is atypical for this early gestational age. C2 had loss of fetal cardiac activity and V2 had findings consistent with severe hydrops fetalis on US. All three cases were humanely euthanized following C-section. No signs of maternal distress were noted in any of these cases.

Pregnant dams that did not experience fetal demise were humanely euthanized after cesarean delivery at a predetermined endpoint (GD90/60 dpi), and complete necropsies were performed. Representative tissue sections (∼1 cm^3^) collected from both dams and fetuses, including muscle, joints, lymphoid tissues, major organs, nervous system, and reproductive organs, were collected into 1mL TRIzol reagent (Invitrogen) for RNA isolation or fixed in 4% paraformaldehyde for histopathology. An additional section from each tissue was preserved in RNAlater for future studies. Early demise embryos and fetuses were mounted fully, as organs were not sufficiently developed for accurate separation. These were subsequently analyzed via RNAscope.

### 6.12. Phenotypic analysis of peripheral blood mononuclear cells

Flow cytometry was performed to phenotype maternal peripheral blood mononuclear cells (PBMCs). Approximately 3 ml of whole blood was centrifuged at 750 G for 15 minutes and plasma was removed and replaced with an equal amount of PBS. The pelleted cellular sample was vortexed before aliquoting 100 μL into three phenotype staining tubes. The remaining sample was layered over lymphocyte separation medium (Corning) and centrifuged for 40 minutes at 2,000 rpm for peripheral blood mononuclear cell (PBMC) isolation. PBMCs were washed in 1X PBS and frozen in liquid nitrogen at ∼2 million cells/ml in freezing medium containing 10%DMSO/40%FBS/50%RPMI.

For immunophenotyping, 100 μL of whole blood was incubated with 1 mL of ACK red blood cell lysis buffer (ThermoFisher) for 5 minutes. Following lysis, cells were pelleted by centrifugation at 2,000 rpm and washed twice with 1X PBS. For analysis of innate immune populations, cells were stained with fluorophore-conjugated antibodies targeting CD45, CD3, CD8a, CD14, CD16, CD11c, HLA-DR, CD56, CD123, and CD169 (activation marker). Monocytes/macrophages were identified as CD3–/CD20–/CD8–/HLA-DR+, with classical (CD14+/CD16–), intermediate (CD14+/CD16+), and non-classical (CD14–/CD16+) subsets delineated accordingly. Dendritic cells (DCs) were classified as CD3–/CD20–/CD8–/HLA-DR+/CD14–/CD16–, further divided into myeloid DCs (CD11c+/CD123–) and plasmacytoid DCs (CD11c–/CD123+). For T cell characterization, staining included CD45, CD3, CD4, CD8a, CD25, CD28, CD95, CD127, CCR6, CXCR3, and intracellular Ki67 and Granzyme B. Naïve CD4+ and CD8+ T cells were CD28+/CD95–, central memory cells were CD28+/CD95+, and effector memory were CD28–/CD95+. Ki67+ and Granzyme B+ populations were gated relative to each animal’s baseline (day 0). For both panels, CD45+ events were first gated to remove residual RBCs, followed by singlet gating prior to subset identification. Data were acquired on a BD Symphony A5 (BD Pharmingen) and analyzed using FlowJo v10.

### 6.13. Ultrasound imaging studies

Ultrasound evaluations were conducted between gestational days 29 and 92 following protocols approved by the Institutional Animal Care and Use Committee (IACUC) for use in established ZIKV pregnancy models (42, 43, 45). Prior to imaging, animals were sedated with an intramuscular injection of ketamine (10 mg/kg), followed by endotracheal intubation and maintenance on inhaled isoflurane (1–2%) for the duration of the procedure at the ONPRC surgical facility. All scans were performed under general anesthesia after an overnight fast. Animals were positioned in dorsal recumbency, and a trained veterinary technician continuously monitored vital signs. Sonographic data were acquired by a board-certified Maternal-Fetal Medicine specialist (J.O.L.) who was aware of the treatment groups using B- Mode, image-directed pulsed and color Doppler on a GE machine (GE Voluson 730) with a 5- to 9- MHz sector probe. Doppler ultrasound measurements were collected with the lowest high-pass filter level (100 Hz) and an angle of 15 or less between the vessel and the Doppler beam.

### 6.14. Fetal biometry

Standard fetal biometric parameters were assessed for biparietal diameter (BPD), head circumference (HC), abdominal circumference (AC), and femur length (FL), as previously described (52, 64–66). Briefly, cranial measurements were acquired in the trans-thalamic imaging plane. BPD was recorded using outer-to-inner caliper placement, while HC was calculated using the ellipse tool, tracing the outer edge of the skull contour. Abdominal circumference was measured in a transverse abdominal plane with the umbilical vein positioned in the anterior third, at the level of the portal sinus. The ellipse tool was similarly applied to trace the outer margin of the abdomen for AC determination. FL was obtained from the femur nearest the transducer, with the shaft positioned as horizontally as possible; calipers were placed at the diaphyseal endpoints, excluding the trochanter. For fetal growth curve comparisons, historical normative data corresponding to the 10th, 50th, and 90th percentiles were previously calculated using established ONPRC linear regression datasets (52).

### 6.15. Amniotic fluid index

Amniotic fluid volume was evaluated following established protocols and ZIKV NHP models (42, 43, 67). The uterine cavity was sectioned into four quadrants, and in each, the deepest vertical pocket of fluid—unobstructed by fetal parts or umbilical cord—was identified and measured. The amniotic fluid index (AFI) was calculated by summing these four individual measurements.

### 6.16. Uterine and umbilical artery volume blood flow

Doppler velocity waveforms of the uterine artery were captured at its proximal segment, as previously described (52, 64). For each animal, three waveform recordings were averaged to determine the pulsatility index (PI), velocity time integral (VTI), and maternal heart rate (MHR) (14). Vessel diameter was assessed using power Doppler angiography(64), and the cross-sectional area (CSA) was calculated using the formula: CSA = (diameter/2)². Uterine artery volume blood flow (cQUta) was then derived using the equation: cQUta = VTI × CSA × MHR. Umbilical artery Doppler assessments were performed on a free-floating segment of the umbilical cord, with measurements collected only during periods of fetal quiescence, absence of breathing or movement (64).

### 6.17. Histopathology

Tissues collected at necropsy for histological analysis were fixed in 4% paraformaldehyde for 24 h and then transferred to 70% ethanol and kept at 4 °C for 24–72 h. The samples were then paraffin-embedded, and slides were cut from the tissue blocks representing the placenta (2 individual 0.5 cm^3^ samples/cotyledon). The slides were stained with hematoxylin and eosin (H&E) on a Leica ST5020 Autostainer (Weltzlar, Germany). Sections from early demise (n = 3, GD35, GD46, GD48), endpoint (n = 3, GD88, GD90, GD92) and gestational age-matched negative controls (n=1; GD35, n=2, GD50 n = 2; GD90) were examined by a board-certified gynecologic pathologist (T.K.M.) blinded to exposure groups. The histopathological assessment included scoring for the presence or absence of placental infarctions, accelerated villous maturation, villous stromal calcifications, chronic villitis, acute chorioamnionitis, and chronic deciduitis (Table 1). Complete cross-sections of decidual spiral arteries were evaluated for luminal dilation and the presence or absence of vasculitis. Tissue slides were examined using Leica DFV495 light microscopes (Wetzlar, Germany) and digitized with the Zeiss Axio scanner (Oberkochen, Germany) at 40× magnification. The histopathological assessments were also made for brain tissue from endpoint cases (C3, V1, V3) by C.S.L. and fetal histopathological assessments were performed by (T.K.M.) unblinded to look for specific pathology related to hydrops fetalis mechanisms.

### 6.18. Immunohistochemistry

Immunohistochemistry was performed as described previously (68). Heat-induced antigen epitope retrieval was performed by heating sections in 0.01% citraconic anhydride containing 0.05% Tween-20 in a pressure cooker (Biocare Medical) set at 110 °C for 15 mins. Slides were incubated with blocking buffer (5% Casein in TBS-T) and then with mouse anti-CD68 monoclonal antibody (1:100), rabbit anti-CD31 monoclonal antibody (1:100), or mouse anti-CD163 monoclonal antibody (1:100) diluted in blocking buffer overnight at room temperature. After washing, endogenous peroxidases were blocked using 1.5% (v/v) H2O2 in PBS, pH 7.4, and the slides were incubated with rabbit or mouse Polink (−1 or −2) horseradish peroxidase and developed with ImmPACT 3,3′- diaminobenzidine (Vector Laboratories), according to the manufacturer’s recommendations. All slides were washed in tap water, counterstained with hematoxylin, mounted in Permount (Thermo Fisher Scientific) and scanned at high magnification (40×) using Zeiss Axio scanner (Oberkochen, Germany).

### 6.19. ZIKV RNA in situ hybridization

ISH was conducted as previously described (69) with the following modifications: Heat-induced epitope retrieval was performed using 1*×* AR9 buffer (Akoya Biosciences cat# AR9001KT) in a Biocare NxGen Decloaker at 110°C for 15 min. Prior to probe application, slides were incubated with diluted (1:10) RNAscope Protease III (ACD cat#: 322337) for 20 minutes at 40°C. Pre-warmed RNAscope Probe ZIKVAsian (ACD cat#: 468361) was incubated on slides for 2 hours at 40°C. Amplification was performed with RNAscope 2.5 HD Reagent Kit RED (ACD cat#: 322350) and washes between amplification steps were performed using RNAscope Wash Buffer (ACD cat#: 320058) used at 0.5*×* concentration. Chromogenic detection after Amplification 6 was performed using Warp Red Chromogen Kit (Biocare cat#: WR806) with a chromogen incubation time of 8 minutes. Slides were counterstained using hematoxylin and lithium bicarbonate blue. An application of Clear-Mount (Electron Microscopy Sciences) was allowed to air-dry before slides were mounted in Permount (Thermo Fisher Scientific).

### 6.20. HALO Integrated Analysis

All slides were scanned at 40*×* magnification on a Zeiss Axio scanner (Oberkochen, Germany). Five vRNA-stained sections per tissue per animal were quantified. Quantitative image analysis was performed in HALO software (Indica Labs-ISH model, v4.2.3* and HALO AI v4.1.5944.161). Organs and compartments were manually annotated from the center slide to delineate major organ systems and compartments for animal C2F. All other animals were evaluated on whole sections, since only sporadic RNA was detected. Uninfected cells were quantified in all annotation layers using Indica Labs – ISH v4.2.3* while ZIKV vRNA+ cells were quantified using a DenseNet v2 HALO AI classifier. The exported data were processed and graphed GraphPad Prism (v.10).

## Author Contributions

All authors have read and agreed to the published version of the manuscript. **Hannah K. Jaeger** Conceptualization, Data curation, Formal analysis, Investigation, Methodology, Validation, Writing – original draft.

**Jessica L. Smith** Conceptualization, Data curation, Formal analysis, Investigation, Methodology, Validation, Writing – original draft,

**Caralyn S. Labriola** Data curation, Formal analysis, Investigation, Methodology, Validation, Writing – review & editing.

**Lydia J. Pung** Data curation, Formal analysis, Investigation, Methodology, Validation, Writing – review & editing.

**Olivia L. Hagen** Data curation, Formal analysis, Investigation, Methodology, Validation, Writing – review & editing.

**Michael Denton** Data curation, Formal analysis, Investigation, Methodology, Validation, Writing – review & editing.

**Rahul J. D’Mello** Data curation, Investigation, Methodology, Writing – review & editing **Christopher J. Parkins** Data curation, Formal analysis, Investigation, Methodology, Validation. **Whitney C. Weber** Data curation, Investigation, Methodology, Writing – review & editing.

**Samuel Medica** Data curation, Investigation, Methodology, Writing – review & editing. **Craig N. Kreklywich** Data curation, Formal analysis, Investigation, Methodology, Validation. **Victor R. DeFilippis** Data curation, Formal analysis, Investigation, Methodology, Validation.

**Stephen Bondoc** Data curation, Formal analysis, Investigation, Methodology, Validation, Writing – review & editing.

**Kathleen Busman-Sahay** Formal analysis, Investigation, Methodology, Validation, Writing – review & editing.

**Jenna N. Castro** Data curation, Formal analysis, Investigation, Methodology, Writing – review & editing.

**Gavin Zilverberg** Data curation, Formal analysis, Investigation, Methodology, Writing – review & editing.

**Riely White** Data curation, Formal analysis, Investigation, Methodology, Writing – review & editing.

**Margaret Terry** Data curation, Formal analysis, Investigation, Methodology, Writing – review & editing.

**Aaron Barber-Axthelm** Data curation, Investigation, Methodology, Writing – review & editing. **Michael K. Axthelm** Data curation, Formal analysis, Investigation, Methodology, Validation, Writing – review & editing.

**Jeremy Smedley** Data curation, Investigation, Methodology, Writing – review & editing. **Matthew T. Aliota** Conceptualization, Data curation, Formal analysis, Methodology, Validation, Writing – review & editing.

**Andrea M. Weiler** Conceptualization, Data curation, Formal analysis, Methodology, Validation, Writing – review & editing.

**Thomas C. Friedrich** Conceptualization, Data curation, Formal analysis, Methodology, Validation, Writing – review & editing.

**Jacob Estes** Conceptualization, Data curation, Formal analysis, Methodology, Validation, Writing – review & editing.

**Terry K. Morgan** Conceptualization, Data curation, Formal analysis, Methodology, Validation, Writing – review & editing.

**Jamie O. Lo** Conceptualization, Data curation, Formal analysis, Investigation, Methodology, Validation, Writing – review & editing.

**Victoria H.J. Roberts** Conceptualization, Data curation, Formal analysis, Investigation, Methodology, Validation, Writing – original draft.

**Daniel N. Streblow** Conceptualization, Data curation, Formal analysis, Investigation, Methodology, Validation, Writing – original draft.

**Alec J. Hirsch** Conceptualization, Data curation, Formal analysis, Investigation, Methodology, Validation, Writing – original draft.

## Institutional Review Board Statement

The study protocol was approved by the Institutional Animal Care and Use Committee of Oregon Health & Science University (protocol code: 0993; expiration date: 12 April 2026).

## Data Availability Statement

The original contributions presented in the study are included in the article/supplementary material; further inquiries can be directed to the corresponding author.

## Acknowledgments

The authors acknowledge the Integrated Pathology Core at the Oregon National Primate Research Center (ONPRC) for the preparation of histologic slides. We would also like to thank the veterinarians and husbandry staff, and the Infectious Disease Resource at the ONPRC who provided excellent care for the animals used in this study.

